# A single chromosome 3p break initiates clear cell renal cell carcinoma evolution

**DOI:** 10.64898/2026.07.10.737056

**Authors:** Rashmi Dahiya, Ianthe A.E.M. van Belzen, Chengheng Liao, Angad Kumar, Yu-Fen Lin, Ahyeon Ko, Amanda K. Mennie, Emilija Aleksandrovic, Jin Zhou, Qing Hu, Justin L. Engel, Jeffrey Miyata, Jose Espejo Valle-Inclán, Alistair G. Rust, Ravikanth Maddipati, James Brugarolas, Srinivas Malladi, Siyuan Zhang, Payal Kapur, Qing Zhang, Isidro Cortés-Ciriano, Peter Ly

## Abstract

Clear cell renal cell carcinoma (ccRCC) is initiated by chromosome 3p loss, yet chromosome losses impose a profound fitness burden on normal cells. How renal epithelial cells tolerate this deleterious aneuploidy during early tumorigenesis remains unclear. Analysis of 949 ccRCC genomes reveals two major classes of chromosome 3p alterations: simple deletions and complex rearrangements surrounding a terminal breakpoint – a pattern we term breakpoint-confined chromothripsis. We modeled both alterations in non-transformed human renal proximal tubule epithelial cells by introducing a single DNA double-strand break on chromosome 3p. Despite an initial fitness disadvantage, chromosome 3p loss drives adaptive genomic evolution that recapitulates recurrent ccRCC-associated aneuploidies, including 5q gain and 14q loss. These alterations alleviate the fitness constraints of 3p loss and promote metabolic reprogramming, clonal expansion, and malignant transformation, producing tumors with features of ccRCC. Thus, a single chromosome break initiates the evolutionary trajectory of ccRCC by creating a fitness bottleneck that selects for recurrent aneuploidies.

## INTRODUCTION

Aneuploidy, defined as the copy number gains and losses of whole chromosomes or chromosome arms, is detected in ∼90% of solid human cancers^1–4^. However, aneuploidy generally imposes a substantial fitness burden through the induction of genotoxic, proteotoxic, and metabolic stress^5–9^. Chromosome losses resulting in partial or complete monosomy are particularly detrimental compared to chromosome gains^10^. These observations raise a longstanding paradox: how can a deleterious chromosomal imbalance become the initiating event during the earliest stages of tumorigenesis? We sought to address this paradox in the setting of clear cell renal cell carcinoma (ccRCC) – the most common histological subtype of kidney cancer.

ccRCC likely originates from the proximal tubules of the kidney^11,12^ and follows a deterministic evolutionary trajectory beginning with loss of the short arm of chromosome 3 (3p) during adolescence – a near-universal genetic hallmark of ccRCC^13–17^. Loss of 3p is followed by expansion of only a few hundred cells^16^, suggesting that rare lineages successfully adapt to this otherwise deleterious genomic state. In sporadic ccRCC, 3p loss represents the ‘first hit’, resulting in the simultaneous heterozygous loss of four tumor suppressor genes located on chromosome 3p: *VHL, PBRM1, BAP1,* and *SETD2*^18^. The *VHL* gene encodes the von Hippel–Lindau tumor suppressor protein (pVHL), a key regulator of cellular oxygen sensing that targets the hypoxia-inducible factor α (HIFα) subunit for degradation^19^. Decades later, inactivating mutations affecting the remaining *VHL* allele, together with one or more of the remaining chromosome 3p tumor suppressor genes, drive oncogenic transformation^16^. Conversely, patients carrying a germline heterozygous *VHL* mutation inevitably undergo 3p loss as the second hit^20,21^. A subset of 3p losses also appears to occur concurrently with a translocation involving a gained copy of chromosome 5q^13,16^. Beyond 3p loss, the gain of 5q and loss of chromosome 14 (commonly referred to as chromosome 14q loss due to its acrocentric structure) represent the second and third most frequent genomic alterations in ccRCC, respectively^22–24^. How these recurrent aneuploidies arise during early cancer development and contribute to malignant progression remains unclear.

DNA double-strand breaks (DSBs) are a major source of genomic instability. Unrepaired DSBs can result in loss of acentric chromosome arms that frequently mis-segregate into micronuclei during mitosis^25^. DSBs can also generate dicentric chromosomes through sister chromatid or non-homologous chromosome fusions. When attached to both spindle poles, dicentric chromosomes form chromatin bridges that may break during anaphase or persist into the subsequent interphase before undergoing fragmentation^26–28^. While chromosome 3p loss in ccRCC has long been considered a simple deletion event, a substantial fraction of tumors harbors localized clusters of rearrangements^16,29^. Such patterns can arise through chromothripsis, a process in which individual chromosomes undergo catastrophic fragmentation followed by error-prone DNA repair^29–34^. Notably, both micronuclei and chromatin bridges can trigger chromothripsis through distinct mechanisms^27,28,31,32^.

Together, these observations raise several key questions. How does chromosome 3p loss arise during early ccRCC evolution? How do cells tolerate the severe fitness costs imposed by this aneuploidy? Lastly, are recurrent alterations such as chromosome 5q gain and 14q loss deterministic adaptations or stochastic consequences of genomic instability? Catastrophic genomic events such as chromothripsis have been proposed as a mechanism underlying concurrent 3p loss and 5q gain in ccRCC^16^, yet whether chromothripsis is required for generating these alterations, or whether stepwise acquisition of DNA copy number changes can similarly drive tumorigenesis, remains unresolved. Moreover, it is unknown whether chromosome 3p loss alone is sufficient to initiate the evolutionary trajectory of ccRCC.

Addressing these questions has been challenging for two reasons. First, the four major tumor suppressor genes located on human chromosome 3p reside on three separate chromosomes in the mouse genome, limiting the ability of genetically engineered mouse models to faithfully recapitulate the genetics of human ccRCC^35^. Second, the severe fitness costs imposed by aneuploidy have hindered efforts to model early cancer-initiating chromosome gains and losses in normal human cells. Consequently, how cells tolerate and adapt to chromosome 3p loss, and how this event primes cells for subsequent malignant progression, remain a central gap in our understanding of ccRCC biology.

Here we identify and characterize a recurrent rearrangement pattern underlying chromosome 3p loss in ccRCC genomes that we term breakpoint-confined chromothripsis. We further engineer an experimental human cell-based model of chromosome 3p loss occurring via breakpoint-confined chromothripsis in non-transformed renal proximal tubule epithelial cells.

Using this system, we show that chromosome 3p loss imposes a severe fitness burden that is overcome through recurrent acquisition of chromosome 5q gain and 14q loss. These alterations drive clonal expansion and malignant progression, providing direct evidence that a deleterious initiating aneuploidy can promote renal tumorigenesis through evolutionary adaptation. More broadly, our findings suggest that similar evolutionary principles may be operative across cancers in which deleterious genetic alterations serve as initiating events.

## RESULTS

### Breakpoint-confined chromothripsis is a recurrent genomic signature of chromosome 3p loss in ccRCC

To investigate the rearrangement mechanisms underpinning chromosome 3p loss in ccRCC, we analyzed whole-genome sequencing (WGS) data from 949 ccRCC tumors from Senkin et al. and The Cancer Genome Atlas (TCGA)^36,37^ **(Methods)**. Consistent with previous studies^38,39^, biallelic inactivation of *VHL* occurred in 76% (723/949) of ccRCC tumors compared to 0.3% (18/7830) of non-ccRCC tumors from TCGA **(Figure S1A-B)**. As expected, loss of chromosome 3p was the most common somatic copy number alteration (SCNA) in ccRCC (93%, 882/949), followed by gain of chromosome 5q and loss of 14q **(Figure 1A)**. The extent of chromosome 3p loss varied across ccRCC tumors, ranging from terminal deletions extending to the *VHL* locus to losses encompassing the entire chromosome arm beyond the centromere **(Figure 1A)**. In 44% (413/949) of ccRCC tumors, the rearrangements underlying chromosome 3p loss harbored the hallmarks of chromothripsis^29^, including clustered breakpoints and a balanced distribution of structural variant (SV) types **(Figure 1B-D, Methods)**. Notably, these rearrangements were typically confined to a small genomic region adjacent to the terminal chromosome 3p breakpoint, a pattern we term ‘breakpoint-confined chromothripsis’.

**Figure 1.**
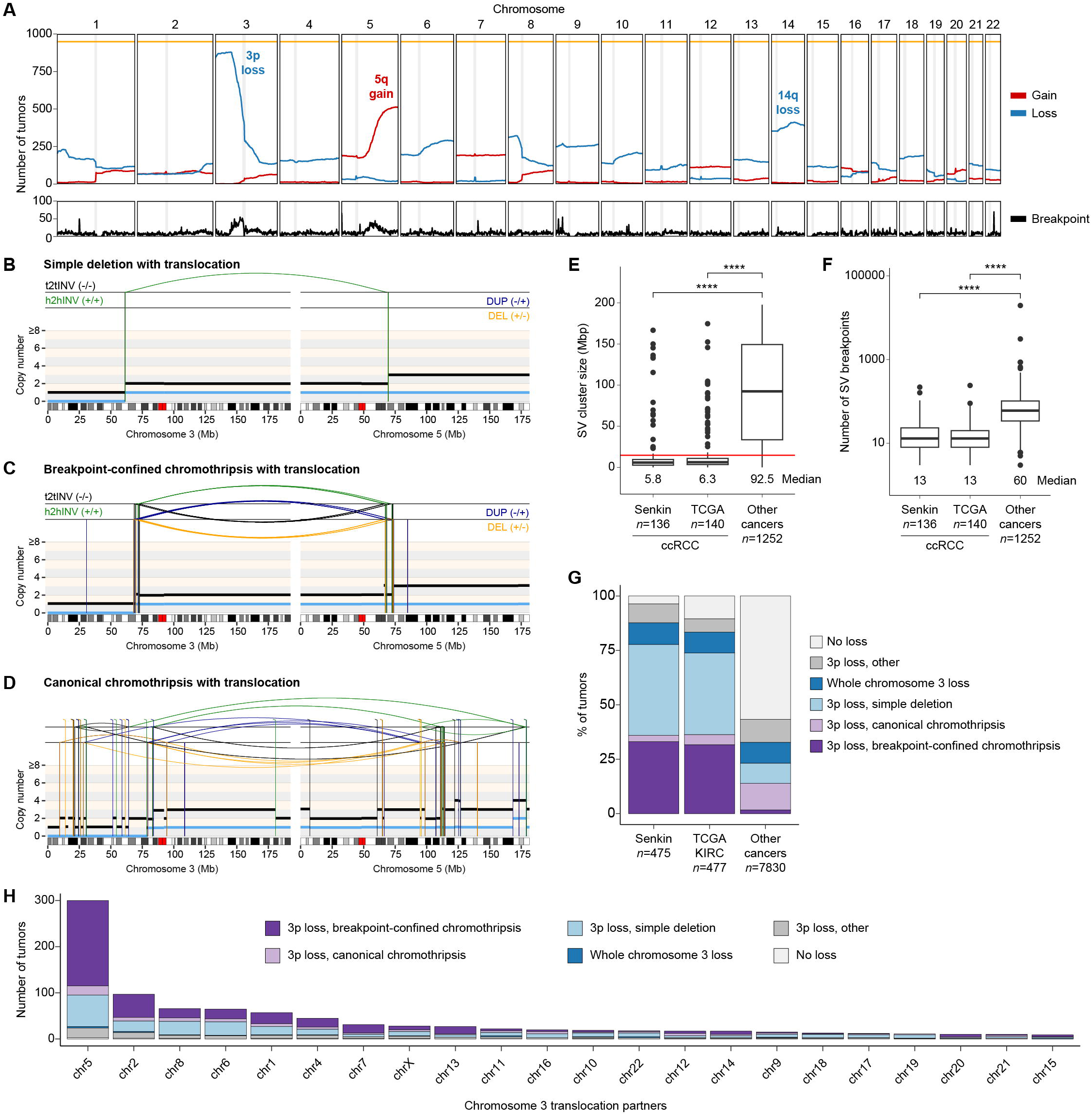
Breakpoint-confined chromothripsis is a recurrent signature of chromosome 3p loss in ccRCC. **(A)** Number of ccRCC tumors with deletions (blue) or gains (red) relative to the tumor ploidy (2n or 4n) based on weighted average copy number across 1 Mbp genomic bins. Breakpoint density is shown in the bottom panel. Centromere positions are indicated by the shaded grey line. **(B-D)** Genomic rearrangement profiles showing with SCNAs (horizontal lines) and SVs (vertical lines) of representative ccRCC tumors with loss of chromosome 3p: (B) simple deletion with translocation involving chromosomes 3 and 5; (C) translocation with breakpoint-confined chromothripsis hallmarks affecting a small genomic region surrounding a terminal breakpoint; **(D)** translocation with canonical chromothripsis hallmarks distributed across both chromosomes. DEL, deletion-like rearrangement; DUP, duplication-like rearrangement; h2hINV, head-to-head inversion; t2tINV, tail-to-tail inversion. **(E)** Size of the genomic region affected by SV clusters detected in tumors with chromosome 3p loss. The changepoint at 14.7 Mbp (red) was used to define breakpoint-confined chromothripsis. *P*<0.01, Wilcoxon rank-sum test. **(F)** Number of chromosome 3 breakpoints for tumors with chromosome 3p loss showing chromothripsis hallmarks. *P*<0.01, Wilcoxon rank-sum test. **(G)** Genomic rearrangements mediating chromosome 3 loss. Breakpoint-confined chromothripsis is defined by SV clusters spanning less than 14.7 Mbp on chromosome 3p and associated with terminal chromosome 3p loss (**Methods**). **(H)** Number of tumors harboring translocations between chromosome 3 and partner chromosomes colored according on the genomic rearrangement underlying chromosome 3p loss.

To determine whether breakpoint-confined chromothripsis is specific to ccRCC, we analyzed WGS data from 7,830 pan-cancer genomes in TCGA. Overall, 44% (3390/7830) of cancers across TCGA harbored chromosome 3p loss, 37% (1252/3390) of which contained SV clusters consistent with chromothripsis **(Figure S1C, Methods)**. However, SV clusters on chromosome 3p in ccRCC (n=276 tumors) spanned substantially smaller genomic regions than those observed in other cancer types (median size of 6.1 versus 92 Mbp, respectively; **Figure 1E**) and involved fewer breakpoints (median 13 versus 60 SVs; **Figure 1F**). Furthermore, chromothripsis events in ccRCC were tightly localized to the most proximal breakpoint of the terminal chromosome 3p loss (median distance of 65 kbp; interquartile range 0-2.6 Mbp), whereas the distance between chromothripsis and chromosome 3p loss was considerably more variable in non-ccRCC tumors (median distance of 1.7 Mbp, interquartile range 0-31 Mbp). Together, these analyses define breakpoint-confined chromothripsis as a recurrent rearrangement pattern characterized by clustered breakpoints spanning less than 14.7 Mbp adjacent to a terminal chromosome 3p loss (indicated by a red line in **Figure 4E, Methods**). Approximately one-third (35%, 307/882) of chromosome 3p loss events were associated with breakpoint-confined chromothripsis in ccRCC compared to 3.8% (130/3390) across all cancers, indicating that this rearrangement pattern is highly enriched in ccRCC evolution **(Figures 1G, S1C)**.

We next explored the mutational mechanisms that could generate breakpoint-confined chromothripsis. Because these events frequently involve two or more chromosomes (94%, 288/307), the presence of multiple translocations together with terminal losses and gains suggests the formation of a derivative chromosome following chromosome breakage and repair. Consistent with this model, translocations involving chromosome 3 were common in tumors with chromosome 3p loss and frequently involved chromosomes with arm-level gains or losses. In agreement with previous studies^16,36^, concurrent loss of 3p and gain of 5q represented the most common combination of alterations in ccRCC (39%, 369/949), approximately half of which were associated with breakpoint-confined chromothripsis (47%, 175/369). Among tumors with chromosome 3 translocations associated with 3p loss (72%, 637/882), translocation partners spanned essentially all human chromosomes but occurred most frequently with chromosome 5 (47%, 297/637), and breakpoint-confined chromothripsis accounted for a substantial fraction of translocations **(Figures 1H, S1D)**. Because detection of chromothripsis requires retention and repair of a sufficient number of genomic fragments, events associated with fragment losses may be indistinguishable from simple translocations. Thus, the prevalence of breakpoint-confined chromothripsis is likely underestimated.

Collectively, these analyses identify breakpoint-confined chromothripsis as a recurrent and highly ccRCC-specific signature of chromosome 3p loss. This rearrangement pattern generates localized clusters of breakpoints adjacent to terminal chromosome losses and frequently produces derivative chromosomes coupling chromosome 3p loss with 5q gain. Thus, breakpoint-confined chromothripsis provides a plausible mechanistic explanation for two of the earliest and most recurrent genomic alterations in ccRCC^16,39^.

### Engineering chromosome 3p loss in renal proximal tubule epithelial cells

To investigate how distinct patterns of chromosome 3p alterations arise and contribute to ccRCC, we engineered a DSB using CRISPR-Cas9 targeted to chromosome 3p in *hTERT*-immortalized, non-transformed human renal proximal tubule epithelial cells, the putative cell of origin of ccRCC^12,40^. To improve genome editing efficiency and facilitate downstream clonal isolation, we expressed CDK4 to accelerate their intrinsically slow proliferation rate, reducing the doubling time from ∼57 to ∼34 hours **(Figure S2A)**. Renal proximal tubule epithelial cells expressing CDK4 (hereafter referred to as RPTECs) displayed enhanced single-cell growth **(Figure S2B)** while maintaining a diploid karyotype without detectable structural abnormalities, as determined by multiplex DNA fluorescent in situ hybridization (FISH) **(Figure S2C)**. Consistent with a stable genomic background, RPTECs exhibited exceptionally low levels of spontaneous micronuclei formation (∼0.5% of cells) **(Figure S2D)**. Inhibition of the mitotic spindle assembly checkpoint increased micronucleation frequency by ∼40-fold **(Figure S2D)**, indicating intact chromosome segregation fidelity under baseline conditions and no evidence of preferential mis-segregation of individual chromosomes.

CRISPR-Cas9-induced DSBs can generate large chromosomal deletions and/or mitotic errors that give rise to abnormal nuclear structures, including micronuclei and chromatin bridges, which are known to provoke chromothripsis^25,41,42^. We hypothesized that an unrepaired acentric 3p fragment would mis-segregate into micronuclei, whereas the centromere-containing chromosome 3 arm may undergo sister chromatid fusion to form dicentric chromosomes, potentially generating mitotic chromatin bridges **(Figure 2A)**. Two Cas9 target sites were selected on 3p near recurrent breakpoint regions observed in ccRCC to encompass all four major 3p tumor suppressor genes **(Figure 2A)**. Delivery of Cas9-sgRNA ribonucleoprotein (RNP) complexes induced micronuclei at a frequency of 3-4%, which increased to 10-15% upon pharmacological inhibition of non-homologous end joining (NHEJ) repair **(Figures 2B-C, S2E)**. DNA FISH confirmed that >70% of micronuclei contained chromosome 3 **(Figure 2D)**, the majority of which were acentric **(Figure S2F)**. Similar results were obtained using lentiviral Cas9 and sgRNA expression, supporting on-target DSB induction at the targeted 3p locus **(Figure S2G-I)**. In parallel, chromosome 3-specific chromatin bridges were observed in 5.2% (sg3p-1, n=293 cells) and 1.6% (sg3p-2, n=197 cells) of anaphase cells **(Figures 2E, S2J)**, and a comparable fraction of interphase cells (∼5%, n=356 cells) harbored persistent chromatin bridges stretched between daughter nuclei **(Figure 2F)**. Thus, targeted DSB induction on chromosome 3p simultaneously generates micronuclei and chromatin bridges in otherwise genomically stable human RPTECs.

**Figure 2.**
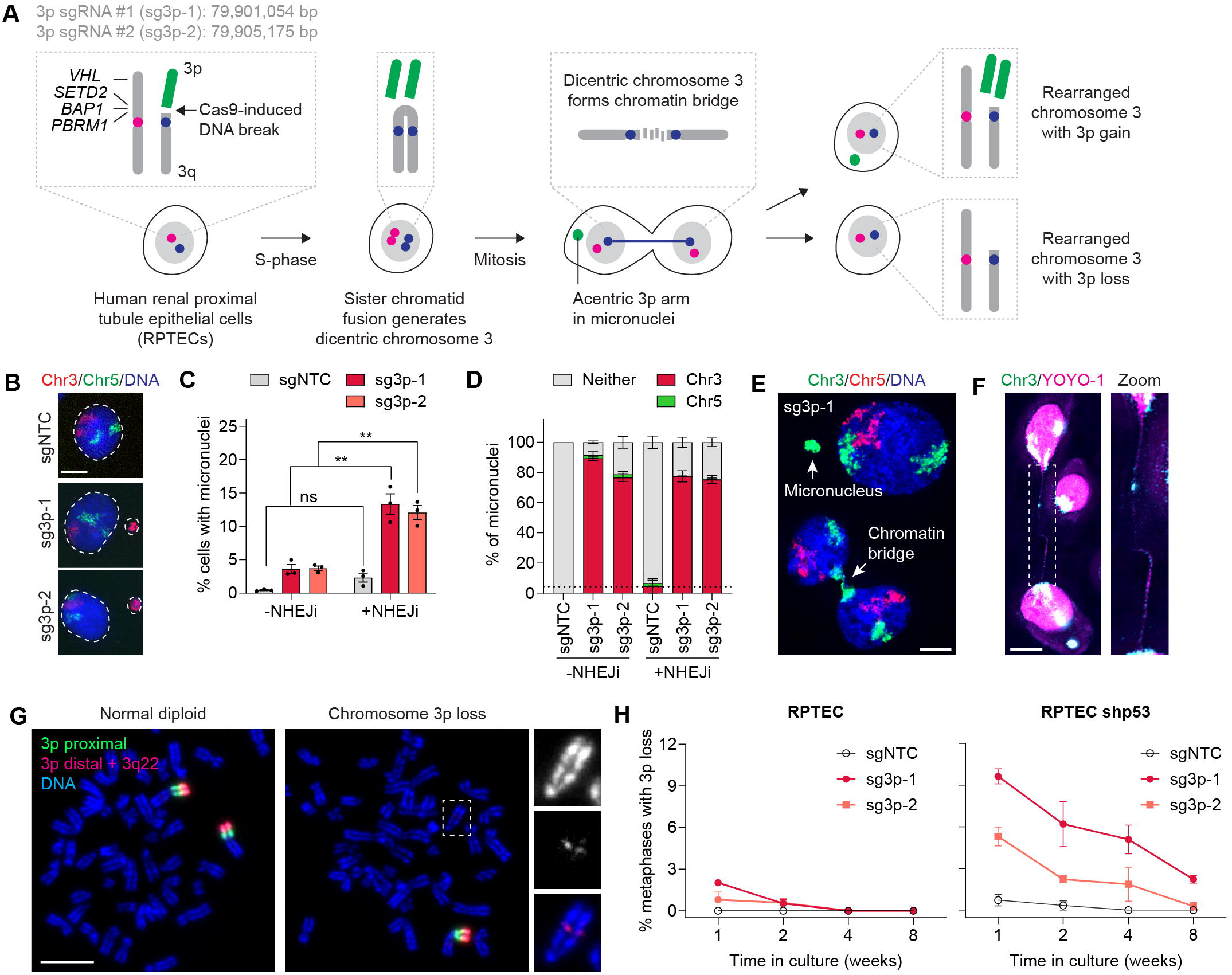
A single chromosome 3p break generates chromosome 3p alterations that are strongly selected against. **(A)** Schematic of the CRISPR-Cas9 strategy used to induce chromosome 3p alterations in non-transformed human renal proximal tubule epithelial cells (RPTECs). Unrepaired acentric fragments mis-segregate into micronuclei, whereas sister chromatid fusion generates dicentric chromosomes that form chromatin bridges. Genomic coordinates of the two chromosome 3p sgRNAs are indicated. **(B)** Interphase FISH images showing chromosome 3-containing micronuclei 72 hr after Cas9-sgRNA RNP delivery. Scale bar, 5 µm. **(C)** Frequency of micronuclei formation following chromosome 3p targeting with or without the DNA-PK inhibitor AZD7648 (NHEJi). Data are mean ± SEM of *n* = 3 experiments from 2984, 1675, 2056, 2129, 1465, 1435, 1438 and 1413 interphase cells (left to right). Statistical analysis by two-way ANOVA test with multiple comparisons. **(D)** Percentage of micronuclei containing chromosome 3 or 5 following chromosome 3p targeting with and without NHEJi. Data are mean ± SEM of *n* = 3 experiments from 14, 58, 72, 92, 32, 192, 173 and 260 micronuclei (left to right). **(E)** Interphase FISH image of a chromosome 3-containing micronucleus and chromosome 3-positive chromatin bridge. Scale bar, 5 µm. **(F)** FISH image of an interphase bridge containing chromosome 3. YOYO-1 is used to label DNA. Right, magnified view of the bridge region. Scale bar, 5 µm. **(G)** Metaphase spreads visualized using dual-colored Oligopaint probes showing a normal chromosome 3 and a chromosome harboring 3p loss. See also **Figure S3**. Scale bar, 10 µm. **(H)** Longitudinal analysis of chromosome 3p loss following Cas9-induced breakage in RPTECs expressing shp53. Data are mean ± SEM of *n* = 3 experiments from 50-280 metaphase spreads per condition at each time point.

To define chromosome 3p-specific alterations arising from Cas9-induced DSBs, we developed a dual-colored Oligopaint strategy leveraging three computationally designed sets of FISH probes. The first probe set hybridizes to the proximal half of 3p, whereas the second targets the distal half of 3p up to the Cas9 cut site. The third probe set targets the 3q22 locus as a control **(Figure S3A)**. In addition to uncovering simple arm-level losses, this configuration enables detection of complex rearrangements through the co-localization of the two Oligopaint probe sets, as previously described^33,43^. Cytogenetic analyses revealed that DSB induction generated a spectrum of 3p alterations, including simple arm-level deletions and inter-chromosomal translocations, the latter of which can be discriminated by the lack of detectable 3q22 signals **(Figure S3B)**. Despite evidence of chromosome 3p fragmentation, highly complex rearrangement patterns characteristic of canonical chromothripsis were not observed **(Figure S3B)**, suggesting that extensive chromosome fragmentation may be poorly tolerated by non-transformed RPTECs.

Because chromosome loss, fragmentation, and rearrangements can activate p53-dependent cell cycle arrest or apoptosis, we suppressed p53 using a short hairpin RNA (shRNA), which reduced p53 phosphorylation and conferred tolerance to ionizing radiation **(Figure S3C-D)**. Following Cas9-induced DSBs, p53-suppressed RPTECs exhibited an increased frequency of 3p loss and other types of 3p alterations **(Figure S3E)**, indicating that p53 acts as an immediate barrier to the persistence and expansion of cells harboring chromosome 3p loss^44,45^. Despite infrequent *TP53* mutations in ccRCC, alternative mechanisms can attenuate p53 pathway activity^46–50^. Notably, cells with 3p loss were readily detectable at early time points but declined toward baseline levels after eight weeks in culture **(Figure 2H)**. These findings suggest that chromosome 3p loss can transiently persist under impaired p53 signaling, but sustained expansion likely requires additional cooperating alterations. Accordingly, all subsequent experiments were conducted using p53-suppressed RPTECs to provide the permissive context necessary for clonal expansion.

We next asked whether selective pressures relevant to ccRCC could promote expansion of cells with chromosome 3p loss. Since ccRCC is characterized by hypoxia-driven HIF stabilization^51,52^, we induced 3p loss and exposed the heterogeneous cell population to two HIF-stabilizing conditions: growth under hypoxia (1% oxygen) and treatment with the prolyl hydroxylase inhibitor dimethyloxalylglycine (DMOG) **(Figure S4A-C)**. However, cytogenetic analyses revealed no increase in the frequency of 3p loss over time under either condition **(Figure S4D)**. Next, to test whether anchorage-independent growth could select for 3p loss^53,54^, we subjected the bulk cell population (comprising ∼10% of cells with 3p loss) to prolonged growth in soft agar yet did not observe enrichment for cells with 3p loss **(Figure S4E-G)**. Finally, to model the germline ccRCC context, we examined whether *VHL* loss could facilitate the persistence of 3p loss cells. Although *VHL* knockout stabilized HIF2α and produced the expected growth defect^55^ **(Figure S4H-I)**, it neither enriched for 3p loss **(Figure S4J)** nor conferred a measurable proliferative advantage **(Figure S3K)**, indicating that activation of the canonical pVHL–HIF signaling pathway is insufficient to overcome the fitness costs imposed by 3p loss. These findings show that, despite being a near-universal truncal event in ccRCC, chromosome 3p loss is subject to strong negative selection across multiple physiologically relevant conditions, suggesting the existence of intrinsic fitness barriers that limit the persistence and expansion of 3p loss cells.

### Chromosome 3p loss drives oxidative stress-induced senescence

To identify transcriptional responses to chromosome 3p loss, we performed single-cell RNA sequencing (scRNA-seq) on bulk cell populations 5 and 10 days following Cas9-induced breakage of chromosome 3p. As a control, we independently targeted the Y chromosome q-arm. InferCNV analysis, which infers large-scale copy number alterations from gene expression patterns across chromosomal regions, identified 3p loss in 14.4% of cells at day 5, declining to 4.8% by day 10 **(Figure S5A-B)**, consistent with cytogenetic measurements **(Figure 2H)**. On-target chromosome 3p breakage was confirmed by reduced expression of 3p-encoded genes distal to the Cas9-induced breakpoint in cells with 3p loss **(Figure S5C)**. Beyond these expected gene dosage effects, scRNA-seq revealed additional transcriptional changes at both time points **(Figure S5D-E)**. Notably, senescence emerged as one of the strongest transcriptional responses^56^ to chromosome 3p loss **(Figure S6A-B)**, and senescence-associated β-galactosidase (SA-β-gal) staining confirmed an elevated fraction of cells exhibiting the enlarged, flattened morphology characteristic of senescence **(Figure S6C-D)**. Gene set enrichment analysis (GSEA) further revealed coordinated enrichment of senescence, oxidative phosphorylation (OXPHOS), reactive oxygen species (ROS), inflammatory response, and p53 pathway signatures **(Figure S6A-G)**, suggesting engagement of a mitochondrial stress-driven, p53-dependent senescence program.

Disruption of *RB1* bypassed this senescence barrier and permitted expansion of cells with chromosome 3p loss **(Figure S6H-I)**. However, this occurred at the expense of elevated whole-genome duplication (WGD) events **(Figure S6L)**, a known mechanism for buffering the deleterious consequences of chromosome loss^57,58^. Thus, while senescence bypass facilitates persistence of chromosome 3p loss cells, it promotes adaptation through an alternative karyotypic route.

The enrichment of OXPHOS and ROS signatures prompted investigation into the mechanistic basis of mitochondrial stress following 3p loss. Elevated OXPHOS drives metastatic progression in ccRCC^59^, yet biallelic *VHL* loss canonically suppresses mitochondrial respiration through HIF-mediated metabolic reprogramming^60,61^. Here, 3p loss resulted in only partial *VHL* haploinsufficiency without detectable activation of hypoxia signaling **(Figure S6A)**, indicating that the OXPHOS signature arises in a HIF-independent context distinct from the pseudohypoxic state characteristic of established ccRCC^61,62^. Because mitochondrial OXPHOS is a principal source of intracellular ROS^63,64^, we reasoned that elevated respiratory activity drives oxidative stress and thereby links chromosome 3p loss to the senescence program described above, as excess ROS can induce cellular senescence^65^. Consistent with this model, cells with 3p loss exhibited robust upregulation of SOD3 **(Figure S5D-E)**, an extracellular superoxide dismutase that mitigates ROS-mediated damage^66^.

To test whether oxidative stress-induced senescence constitutes a barrier to persistence of 3p loss cells, we generated RPTECs overexpressing SOD3 **(Figure S6J)** and subsequently induced Cas9-mediated chromosome 3p DSBs. SOD3 overexpression enriched for 3p loss cells relative to controls **(Figure S6K)** and, in contrast to *RB1*-mediated senescence bypass, did so without a concomitant increase in WGD **(Figure S6L)**. These data identify oxidative stress-induced senescence as a major fitness barrier imposed by chromosome 3p loss and suggest that redox adaptation represents an early evolutionary solution that permits aneuploidy tolerance.

### dTAG-Flow enables isolation of rare clones with chromosome 3p loss

We next sought to isolate RPTEC clones with chromosome 3p loss for characterization. Initial screening of 162 single cell-derived clones by FISH failed to identify any clones with 3p loss. To enrich for the rare cells that persist despite 3p loss, we developed a degron-based fluorescence-activated cell sorting (FACS) strategy that leverages the E3 ubiquitin ligase activity of pVHL as a functional surrogate for chromosome 3p status. To do so, we expressed green fluorescent protein (GFP) fused to an FKBP12^F36V^ degron (hereafter referred to as GFP-degron). Upon treatment with the pVHL-mediated heterobifunctional degrader dTAGv1^67^, GFP-degron undergoes proteasomal degradation in cells with intact pVHL. In contrast, cells with 3p loss exhibited reduced pVHL activity and retention of GFP-degron fluorescence, thereby enabling their isolation by FACS. We termed this strategy dTAG-Flow **(Figure 3A)**. Validation in *VHL*-edited RPTECs confirmed that dTAGv1 induced complete degradation of GFP-degron in WT cells **(Figure S7A-E)** but only partial or no degradation in *VHL^+/-^* and *VHL^-/-^*cells, respectively **(Figure S7D-E)**. Thus, GFP-degron retention provides a functional readout of reduced pVHL activity and enables enrichment of cells with chromosome 3p loss.

**Figure 3.**
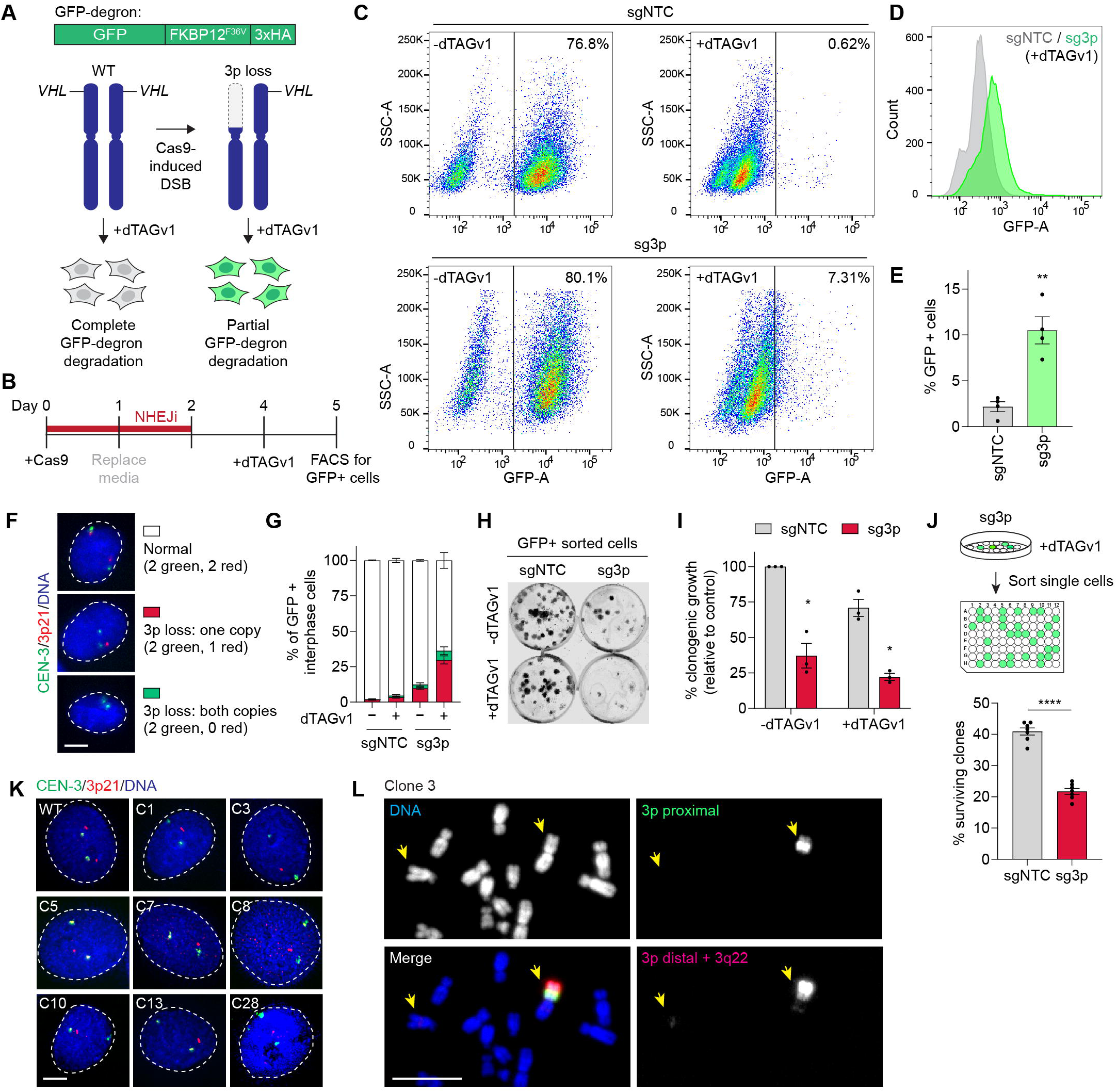
dTAG-Flow enables isolation of rare clones harboring chromosome 3p loss. **(A)** Schematic of the dTAG-Flow strategy for enrichment of cells with reduced pVHL activity following chromosome 3p loss. **(B)** Experimental workflow for dTAG-Flow. **(C-D)** Flow cytometry profiles and quantification of GFP+ cells following chromosome 3p targeting and dTAGv1 treatment. **(E)** Frequency of GFP+ cells recovered by dTAG-Flow. Data are mean ± SEM of *n* = 4 independent experiments. Statistical analysis by one-way ANOVA test with multiple comparisons. **(F)** Interphase FISH images of GFP+ cells immediately following dTAG-Flow enrichment. Scale bar, 5 µm. **(G)** Quantification of chromosome 3p status in GFP+ cells shown in (F). Data are mean ± SEM from *n* = 3 experiments from 411, 342, 333, and 345 interphase cells (left to right). **(H)** Clonogenic survival assay of GFP- and GFP+ populations following dTAG-Flow enrichment. **(I)** Quantification of clonogenic survival shown in (H). Data are mean ± SEM from *n* = 3 experiments. Statistical analysis by two-way ANOVA test with multiple comparisons. **(J)** Frequency of surviving single-cell clones recovered following dTAG-Flow enrichment. Data are mean ± SEM from *n* = 5 experiments. Statistical analysis by paired t-test. **(K)** Interphase FISH images of independently derived clones with chromosome 3p loss isolated by dTAG-Flow. Scale bar, 5 µm. **(L)** Metaphase spread from clone 3 visualized using dual-colored Oligopaint probes showing the loss of a chromosome 3p arm. Scale bar, 10 µm.

To isolate RPTECs with 3p loss, we applied dTAG-Flow following Cas9-induced chromosome 3p breakage **(Figure 3B)**. Targeting 3p resulted in an enrichment of GFP+ cells compared to controls, consistent with impaired degradation of GFP-degron due to reduced pVHL activity **(Figure 3C-E)**. Interphase FISH immediately following sorting confirmed that the bulk GFP+ population was enriched for cells with 3p loss **(Figure 3F-G)**. Consistent with a strong fitness deficit associated with 3p loss, GFP-sorted cells exhibited reduced viability **(Figure 3H-I)** with a corresponding depletion of 3p loss cells over time **(Figure S8A)**.

To capture rare surviving cells, GFP+ cells were single-cell sorted into 96-well plates to derive clonal populations. Despite reduced clonogenic growth **(Figure 3J)**, we expanded 40 surviving clones from two independent experiments and screened them for 3p loss using interphase FISH probes targeting the chromosome 3 centromere and the 3p21 locus **(Figure S8B)**, yielding eight independent RPTEC clones harboring chromosome 3p loss **(Figure 3K)**. These clones were validated on metaphase spreads using the previously described dual-colored Oligopaint probes **(Figure 3L)**, as well as whole-chromosome paint probes for chromosome 3 and a control chromosome 5 **(Figure S8C)**. Thus, dTAG-Flow enables isolation and expansion of rare, genetically defined chromosome 3p loss clones from an otherwise counterselected population.

As expected, bulk RNA sequencing of clones with 3p loss showed reduced expression of the four major ccRCC tumor suppressor genes encoded on chromosome 3p **(Figure S8D)**, consistent with gene dosage effects resulting from hemizygous chromosome loss. Baseline micronuclei formation remained low across clones **(Figure S8E)**, and γH2AX immunostaining revealed variable levels of DNA damage, indicative of low to moderate levels of genomic instability **(Figure S8F)**. FISH analyses further detected telomeric sequences at chromosome 3p breakpoints **(Figure S8G)**, suggesting stabilization of broken chromosome ends and maintenance of derivative chromosome with terminal arm losses. Notably, all 3p loss clones exhibited robust induction of SOD3 **(Figure S8H)**, consistent with the transcriptional response observed by scRNA-seq **(Figure S5D-E)** and supporting a compensatory adaptation to oxidative stress. Although SOD3 is typically localized extracellularly or in the cytoplasm, it accumulated within the nucleus of all 3p loss clones **(Figure S8I-J)**, indicative of altered redox regulation associated with 3p loss^68,69^. These findings indicate that 3p loss cells engage persistent stress response programs that may facilitate survival despite the severe fitness constraints imposed by this aneuploidy. These isogenic models provide a tractable system to interrogate how cells adapt to chromosome losses and how a detrimental initiating lesion shapes subsequent stages of ccRCC evolution.

### Recurrent chromosome 5q gain and 14q loss promotes clonal evolution after chromosome 3p loss

All eight clones with chromosome 3p loss proliferated more slowly than their diploid isogenic counterparts **(Figure 4A)**, consistent with a substantial fitness burden caused by 3p loss. To determine whether 3p loss is sufficient to confer features of malignant transformation, we subjected these clones to anchorage-independent growth assays^70^. Three clones (7,10 and 28) formed robust colonies in soft agar, whereas the remaining clones exhibited minimal or no growth **(Figure 4B-C)**. These results indicate that 3p loss is not uniformly sufficient for malignant growth and that additional subclonal alterations influence tumorigenic potential, consistent with observations in other epithelial systems^71^.

**Figure 4.**
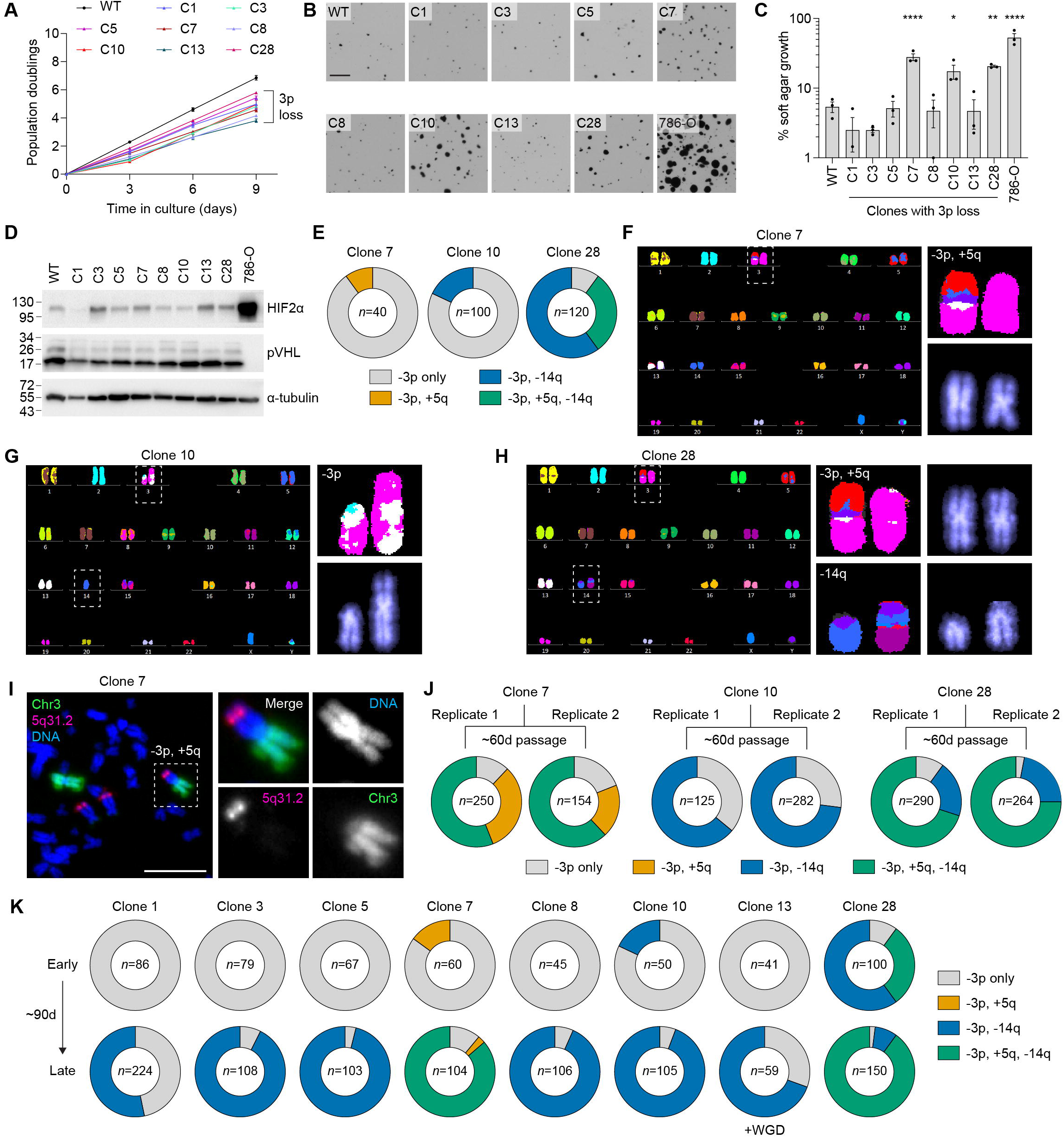
Adaptive aneuploidies drive tumorigenic evolution following chromosome 3p loss. **(A)** Cumulative population doublings of WT RPTECs and chromosome 3p loss clones. **(B)** Images of soft agar colony formation assays of WT RPTECs and chromosome 3p loss clones. Scale bar, 1 mm. **(C)** Quantification of anchorage-independent growth shown in (B). Data are mean ± SEM from *n* = 3 independent experiments. Statistical analysis by two-way ANOVA test with multiple comparisons. **(D)** Immunoblot analysis of HIF2α and VHL expression across chromosome 3p loss clones compared to WT RPTECs. **(E)** Frequency of recurrent ccRCC-associated chromosomal alterations detected in clones 7, 10, and 28. Sample sizes indicate number of karyotypes analyzed by metaphase FISH per clone. **(F)** Representative multiplex FISH karyotype of clone 7 showing a translocated chromosome with chromosome 3p loss and 5q gain. **(G)** Representative multiplex FISH karyotype of clone 10 showing chromosome 14q loss. Because chromosome 14 is acrocentric, whole-chromosome 14 loss is referred to as chromosome 14q loss. **(H)** Representative multiplex FISH karyotype of clone 28 showing a translocated chromosome with chromosome 3p loss and 5q gain along with chromosome 14q loss. **(I)** Chromosome paint analysis of clone 7 using chromosome 3 and 5 paint probes. Insets show the derivative chromosome with a translocation between chromosomes 3p and 5q. Scale bar, 10 µm. **(J)** Longitudinal tracking of recurrent ccRCC-associated aneuploidies following serial passaging of clones 7, 10, and 28 into two separate replicates. Sample sizes indicate number of karyotypes analyzed by metaphase FISH per clone. **(K)** Convergent karyotypic evolution of chromosome 3p loss clones following prolonged culture, characterized by recurrent acquisition of chromosome 5q gain and chromosome 14q loss. Sample sizes indicate number of karyotypes analyzed by metaphase FISH per clone. WGD, whole-genome duplication.

To determine whether variation in malignant potential could be explained by differential activation of the canonical pVHL–HIF2α pathway, we assessed HIF2α levels across the clone panel **(Figure 4D)**. Although half of the 3p loss clones exhibited partial stabilization of HIF2α under normoxic conditions **(Figure 4D)**, HIF2α abundance did not always correlate with anchorage-independent growth. For instance, clones 3 and 13 showed detectable HIF2α stabilization yet failed to form colonies in soft agar **(Figure 4B-C)**. These findings indicate that partial *VHL* haploinsufficiency and modest HIF2α accumulation are insufficient to explain differences in malignant potential.

We next hypothesized that additional genomic alterations cooperate with 3p loss to promote malignant growth. Chromosome painting and multiplex FISH revealed that the tumorigenic clones (7, 10, and 28) acquired subclonal chromosomal alterations that are recurrently observed in ccRCC, whereas non-tumorigenic clones did not **(Figures 4E-I, S9)**. Specifically, clone 7 harbored a translocation between 3p and a gained 5q arm in 10% of cells, clone 10 showed loss of chromosome 14q in 18% of cells, and clone 28 contained both a translocation between 3p and a gained 5q and loss of chromosome 14q in 30% of cells, with chromosome 14q loss alone present in an additional 60% of cells (**Figure 4E**). Because chromosome 14 is acrocentric and its short arm primarily contains ribosomal DNA repeats, whole-chromosome 14 loss detected by cytogenetics is functionally equivalent to the chromosome 14q losses reported in ccRCC genomic studies. Notably, 5q gain and 14q loss represent the two most frequent secondary genomic alterations in ccRCC **(Figure 1A)**^16,24^, indicating that independent 3p loss clones converge toward stereotyped evolutionary trajectories.

To define the genomic landscape of these clones at higher resolution, we performed WGS, which confirmed the subclonal alterations **(Figure S10)** identified by multiplex FISH. Remarkably, two of eight clones (7 and 8) exhibited rearrangement patterns consistent with breakpoint-confined chromothripsis, characterized by localized clusters of SVs surrounding terminal 3p breakpoints. Although less extensive than the patterns observed in human ccRCC tumors, these findings demonstrate that a single chromosome 3p break can generate rearrangement signatures resembling those identified in patient tumors.

To determine whether recurrent ccRCC-associated alterations confer a selective advantage, we serially passaged two independent replicates of clones 7, 10, and 28 for an additional 60 days and longitudinally tracked their karyotypes by metaphase FISH. Strikingly, the frequency of 5q gain increased from ∼10% to ∼70% in clone 7 and from 30% to 75% in clone 28 over the course of passaging **(Figure 4J)**. Similarly, chromosome 14q loss in clone 10 increased from ∼18% to ∼70% by the endpoint **(Figure 4J)**. Notably, all three clones ultimately acquired chromosome 14q loss, revealing convergent evolution toward a ccRCC-like genomic landscape. These findings indicate that 5q gain and chromosome 14q loss confer a selective clonal advantage in the context of 3p loss, driving progressive clonal expansion of cells harboring these recurrent alterations.

We next asked whether restoration of p53 signaling could restrain the growth of tumorigenic 3p loss clones. To do so, we expressed a doxycycline-inducible, shRNA-resistant *TP53* cDNA in clones 7 and 10 **(Figure S11A)**. Induction with doxycycline reduced both cell proliferation and anchorage-independent growth **(Figure S11B-E)**, consistent with our earlier observations that p53 constrains the persistence and expansion of cells with 3p loss **(Figure 2H)**. However, restoration of p53 signaling was insufficient to reverse the underlying karyotypic abnormalities, as assessed by metaphase FISH **(Figure S11F-G)**, indicating that p53 suppresses clonal expansion without correcting established chromosomal alterations. These findings suggest that 3p loss creates a selective environment in which recurrent aneuploidies such as 5q gain and chromosome 14q loss emerge and expand by alleviating the fitness costs imposed by the initiating lesion. Thus, adaptive chromosomal evolution, rather than chromosome 3p loss alone, drives sustained tumorigenic growth.

### Convergent chromosome 14q loss drives metabolic adaptation during early ccRCC evolution

We next serially passaged all 3p loss clones for 90 days while longitudinally monitoring chromosomal alterations by cytogenetics. Remarkably, every clone independently acquired *de novo* chromosome 14q loss in the context of pre-existing 3p loss **(Figure 4K)**. This recurrent event was observed across genetically distinct clones, indicating strong positive selection rather than stochastic accumulation of chromosomal alterations. Notably, clones 7 and 28 subsequently acquired chromosome 14q loss on a background of pre-existing 5q gain involving a 3p translocation, resulting in layered aneuploidy within 3p loss cells **(Figure 4K)**. The sequential acquisition of chromosome 3p loss, 5q gain, and 14q loss closely mirrors the evolutionary trajectory observed in ccRCC. While some early-passage 3p loss clones maintained relatively stable karyotypes, others (clones 7, 10, and 28) progressively acquired subclonal ccRCC-associated alterations that expanded over time **(Figure 4E, 4J)**. However, all eight late-passage clones ultimately developed chromosome 14q loss, reinforcing convergence toward a common evolutionary endpoint **(Figure 4K)**. The universal emergence of chromosome 14q loss across independent clones with 3p loss represents a strong example of convergent karyotypic evolution observed in our system.

To define these evolutionary changes at higher resolution, we compared the genomes of early- and late-passage clones with 3p loss by WGS **(Figures S10, S12)**. Both early- and late-passage clones acquired additional copy number alterations and SVs that recapitulate patterns observed during ccRCC progression **(Figure 5A-D)**. These rearrangements arose in the absence of exogenous mutagenic stress, indicating that the altered genomic state initiated by 3p loss itself is sufficient to drive ongoing genome evolution. Notably, breakpoint-confined chromothripsis of 3p with and without detectable translocations emerged in our model system, recapitulating the spectrum of chromosome 3p alterations observed in patient tumors **(Figures 1, 5A-D)**.

**Figure 5.**
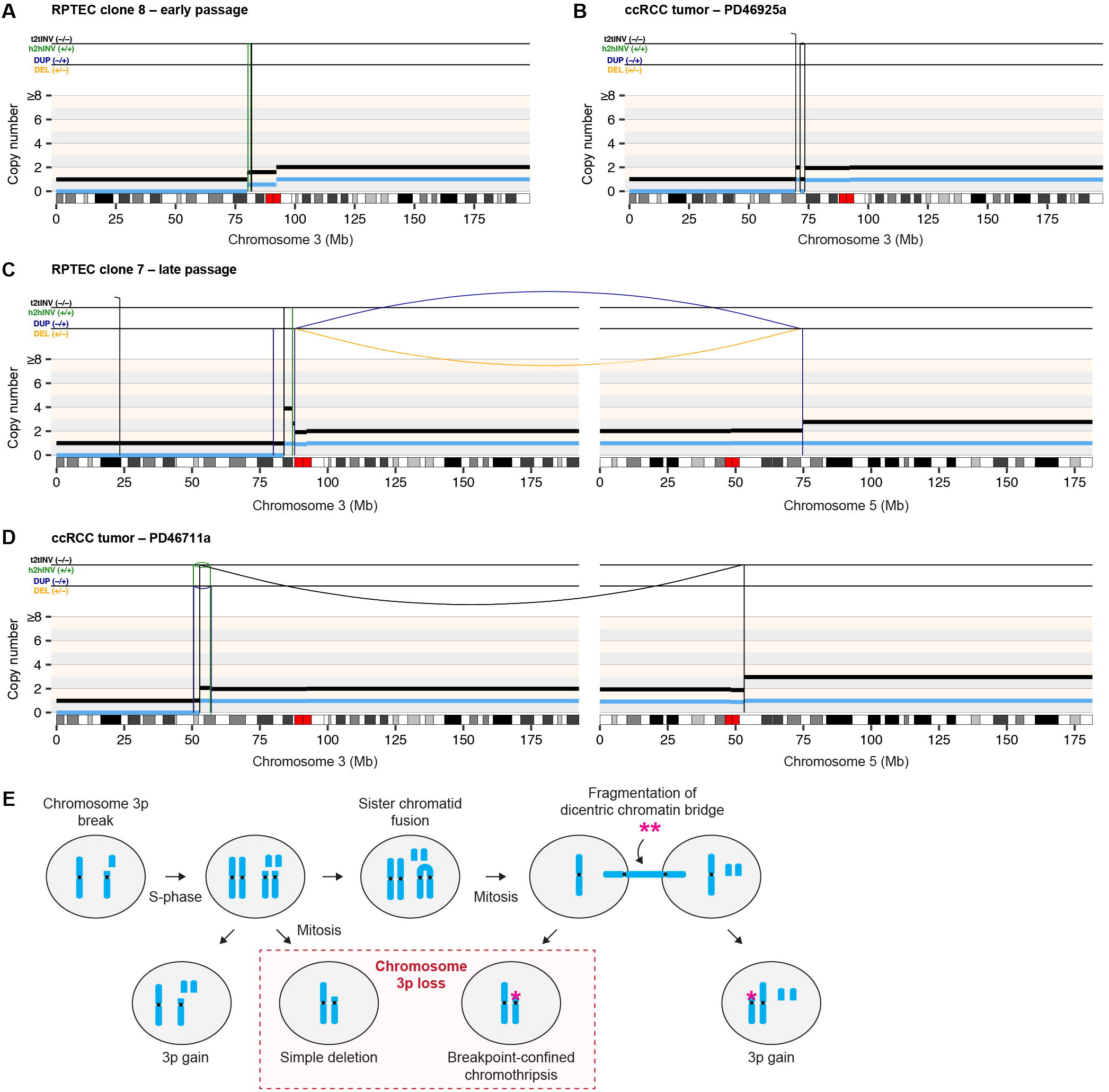
A single chromosome break generates the spectrum of chromosome 3p alterations observed in ccRCC. **(A-B)** Genomic rearrangement profiles showing breakpoint-confined chromothripsis associated with chromosome 3p loss from an early-passage chromosome 3p loss clone (A) and human ccRCC tumor (B). SVs are clustered adjacent to the terminal chromosome 3p breakpoint and are accompanied by localized copy number alterations. **(C-D)** Genomic rearrangement profiles showing concurrent chromosome 3p loss and 5q gain with a translocation from a late-passage chromosome 3p loss clone (C) and human ccRCC tumor (D). Colored arcs indicate SV junctions linking chromosomes 3 and 5. **(E)** Proposed model for generating chromosome 3p alterations following a single chromosome break. Loss of the acentric chromosome 3p fragment generates simple chromosome 3p deletions. Sister chromatid fusion and bridge fragmentation generate breakpoint-confined chromothripsis. Subsequent breakage-fusion-bridge cycles can produce derivative chromosomes coupling chromosome 3p loss with a translocation, which most commonly involves chromosome 5q gain. Asterisks denote clustered SVs.

These observations support a mechanistic model for the origin of recurrent 3p alterations in ccRCC **(Figure 5E)**. Following a DSB on chromosome 3p, the acentric 3p arm can be lost through mitotic mis-segregation, resulting in a simple arm-level deletion. Alternatively, replication of the broken chromosome followed by sister chromatid fusion generates a dicentric chromosome that forms a chromatin bridge during mitosis. Local fragmentation and repair of this bridge produce breakpoint-confined chromothripsis, yielding daughter cells with 3p loss and localized structural rearrangements. In either of the resulting daughter cells, the remaining chromosome 3 end may be stabilized by telomere acquisition or trigger an additional BFB cycle, which can involve chromosome 5 and generate derivative chromosomes harboring concurrent chromosome 3p loss and 5q gain. Thus, a single initiating DSB can give rise to both simple deletions and breakpoint-confined chromothripsis, providing a unified explanation for the earliest recurrent genomic alterations observed in ccRCC.

Because chromosome 14q loss emerged as a recurrent evolutionary endpoint, we next sought to understand its functional consequences. Chromosome 14q loss occurs in over 40% of ccRCC tumors and is associated with advanced stage, higher grade, and poor clinical outcome^22–24,72^ ^73^. A compelling candidate underlying its selective advantage is *HIF1A*, located on chromosome 14q and encoding for HIF1α, which functions as a tumor suppressor in ccRCC. In contrast, *EPAS1* (encoding for HIF2α), located on chromosome 2p and activated upon *VHL* loss, functions as an oncogenic driver^74^. In this context, loss of chromosome 14q may represent an adaptive step that shifts the balance toward pro-tumorigenic signaling. In agreement with this model, chromosome 14q loss conferred a proliferative advantage **(Figure 6A)** in all but one clone (clone 13), which instead underwent WGD **(Figures 4K)**. Moreover, late-passage clones exhibited enhanced anchorage-independent growth, indicative of increased tumorigenic potential **(Figure 6B-C)**. Although endogenous HIF1α levels remained largely unchanged despite hemizygous chromosome 14q loss **(Figure S13A)**, overexpression of HIF1α did not affect two-dimensional growth **(Figure S13B-C)** but partially reduced soft agar colony formation across independent clonal populations **(Figure S13D-E)**, supporting the notion that attenuation of HIF1α activity contributes to the selective advantage associated with chromosome 14q loss. These findings suggest that chromosome 14q loss promotes ccRCC evolution, at least in part, by alleviating HIF1α-mediated growth restraint.

**Figure 6.**
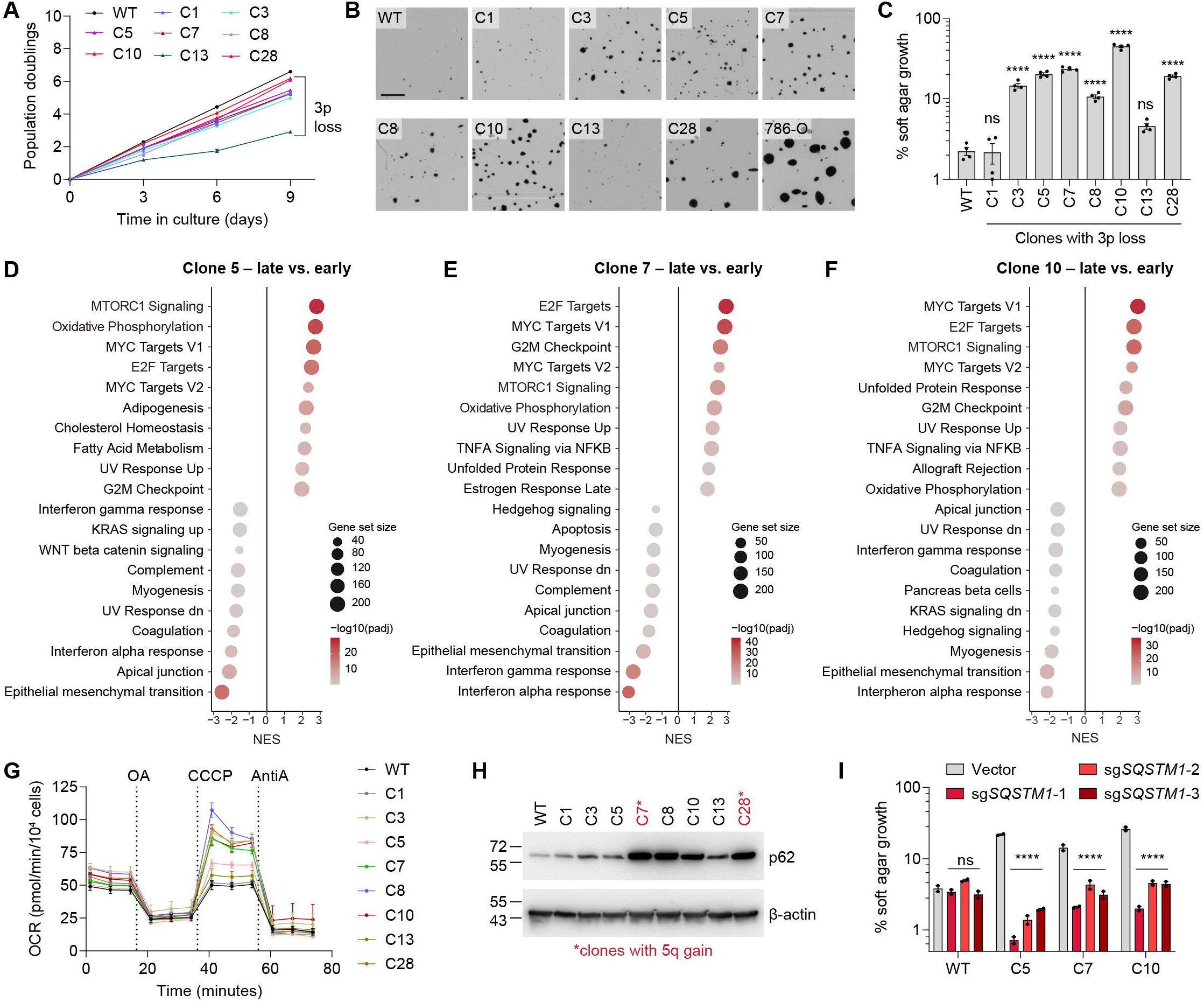
Convergent chromosome 14q loss drives metabolic adaptation during early ccRCC evolution. **(A)** Cumulative population doublings of WT RPTECs and late-passage chromosome 3p loss clones. **(B)** Images of soft agar colony formation assays of late-passage chromosome 3p loss clones and WT RPTECs. Scale bar, 1 mm. **(C)** Quantification of anchorage-independent growth shown in (B). Data are mean ± SEM from *n* = 3 independent experiments. Statistical analysis by two-way ANOVA test with multiple comparisons. **(D-F)** Hallmark pathway enrichment analysis comparing early- and late-passage clones 5, 7, and 10. Positive normalized enrichment scores (NES) indicate pathways enriched during clonal evolution. **(G)** Seahorse analysis showing enhanced mitochondrial respiratory capacity in late-passage chromosome 3p loss clones. OCR, oxygen consumption rate. Data are mean ± SEM from *n* = 8 technical replicates per cell line. **(H)** Immunoblot analysis showing elevated p62 expression in late-passage chromosome 3p loss clones. **(I)** Quantification of anchorage-independent growth following inactivation of SQSTM1 (p62) in the indicated clones (also see **Figure S13J**). Data are mean ± SEM from *n* = 3 independent experiments. Statistical analysis by two-way ANOVA test with multiple comparisons.

To define the molecular consequences of chromosome 14q loss in the setting of pre-existing 3p loss, we compared the transcriptomes of early- and late-passage clones by bulk RNA sequencing. Clones with acquired chromosome 14q loss exhibited a distinct transcriptional program characterized by upregulation of mTORC1 signaling and OXPHOS pathways (**Figures 6D-F, S13F-H**). Consistent with these transcriptional changes, Seahorse analysis revealed markedly increased maximal and spare respiratory capacity, demonstrating enhanced mitochondrial respiratory function **(Figure 6G)**. Thus, acquisition of chromosome 14q loss is associated with metabolic reprogramming and enhanced nutrient-sensing activity, potentially conferring a proliferative and survival advantage consistent with ccRCC progression^75^.

p62 (encoded by *SQSTM1* on chromosome 5q), an established activator of mTORC1^76^ and a proposed pathogenic target of 5q amplification in ccRCC^77^, emerged as a candidate mediator of this phenotype. Surprisingly, p62 protein levels were elevated in 3p loss clones irrespective of chromosome 5q status; for example, clones 7, 8, and 10 exhibited comparable p62 abundance despite differing chromosome 5q copy numbers **(Figure 6H)**. To assess whether p62 is functionally necessary, we deleted *SQSTM1* in clones 5, 7, and 10 using three independent sgRNAs **(Figure S13I)**. *SQSTM1* loss substantially impaired anchorage-independent growth in all three chromosome 3p loss clones **(Figures 6I, S13J)**, identifying p62 as a functionally required mediator of the tumorigenic properties acquired during sequential chromosomal evolution. Recurrent aneuploidies therefore act as adaptive solutions to the fitness constraints imposed by 3p loss, promoting metabolic reprogramming, clonal expansion, and tumorigenic progression.

### Sequential chromosomal evolution culminates in tumorigenesis and metastasis

To determine whether the recurrent chromosomal alterations acquired during long-term evolution were sufficient for malignant transformation *in vivo*, we performed simultaneous subcutaneous injections of early- and late-passage derivatives of clones 7 and 10 alongside diploid RPTECs into immunocompromised mice. Early-passage cells failed to form tumors during ∼400 days of observation, whereas late-passage cells generated xenograft tumors in as little as ∼120 days **(Figures 7A-C, S14A**). Notably, all tumors arising from late-passage clone 10 exhibited spontaneous metastatic dissemination to the lung **(Figure 7B-C)**.

**Figure 7.**
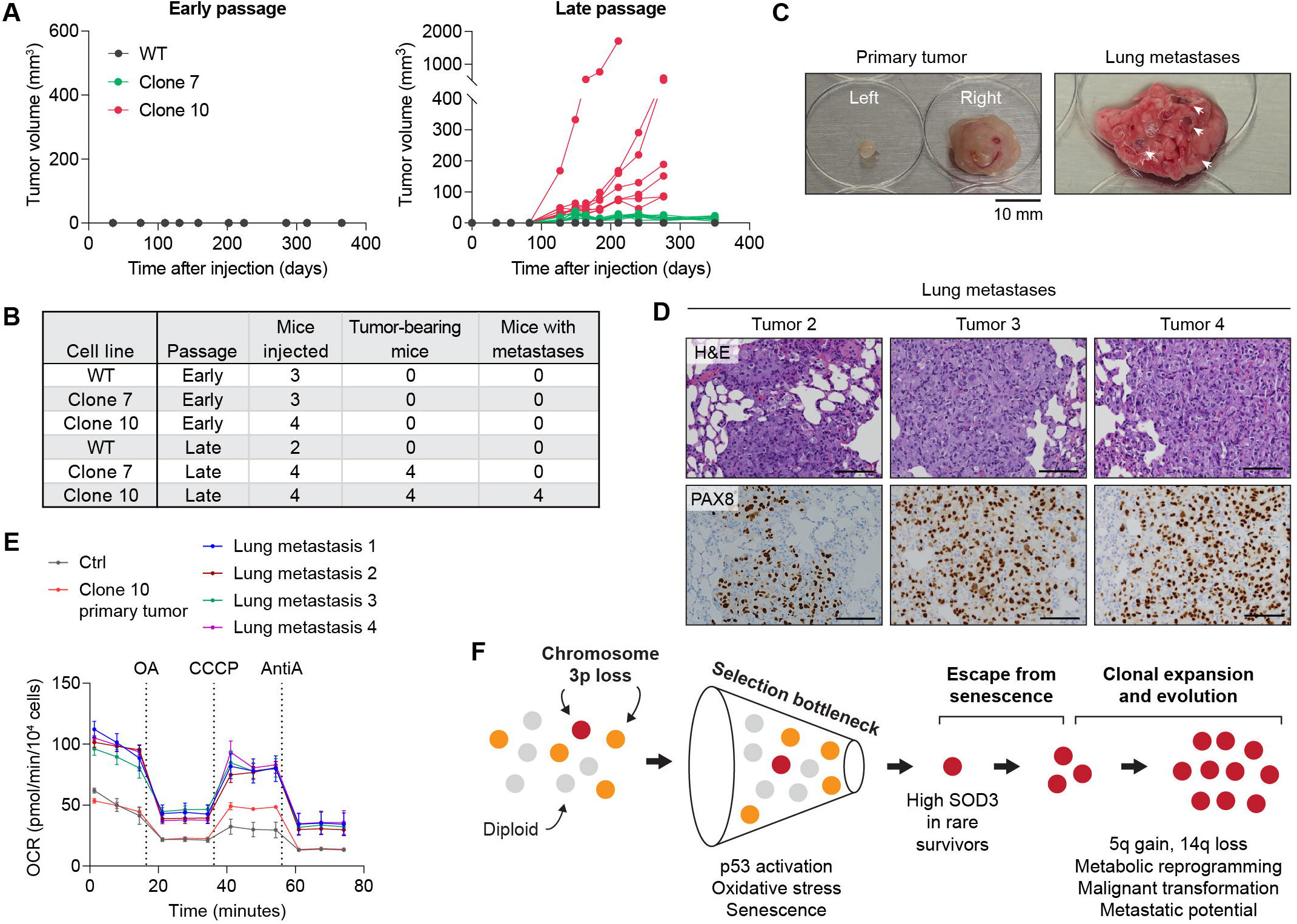
Sequential chromosomal evolution culminates in renal tumorigenesis and metastasis. **(A)** Xenograft growth of early- and late-passage chromosome 3p loss clones and WT RPTECs following subcutaneous implantation into immunocompromised mice. **(B)** Summary of tumor incidence and metastatic dissemination following implantation of early-and late-passage cell populations. **(C)** Images of primary tumor and spontaneous lung metastases arising tumor 1 derived from late-passage clone 10 cells. Arrows indicate metastatic lung nodules. **(D)** Representative hematoxylin and eosin (H&E) staining and immunohistochemistry (IHC) for the renal lineage marker PAX8 in lung metastases derived from late-passage clone 10 tumors. Scale bar, 100 µm. **(E)** Oxygen consumption rate (OCR) measurements of cells isolated from primary tumors and lung metastases compared with WT RPTECs. Data are mean ± SEM from *n* = 8 technical replicates per cell line. **(F)** Proposed model of ccRCC evolution. Chromosome 3p loss creates a fitness bottleneck that selects for recurrent adaptive aneuploidies, including chromosome 5q gain and chromosome 14q loss, driving metabolic reprogramming, malignant growth, and metastatic potential.

Histopathological examination of hematoxylin and eosin-stained sections from primary tumor 1 revealed a malignant epithelial neoplasm composed of cells arranged in nests and sheets with focally clear cytoplasm and separated focally by thin capillary vasculature, imparting focal ccRCC-like morphology **(Figure S14B)**. Tumors 2-4 and the corresponding lung metastases were composed predominantly of solid sheets of neoplastic cells with variable fibrosis, which was particularly prominent in tumors 2 and 4 **(Figures 7D, S14B)**. Immunohistochemically, all tumors showed strong expression of the renal lineage marker PAX8 in both primary tumors and lung metastases **(Figures 7D, S14B)**.

Unexpectedly, immunohistochemical analyses revealed that the primary tumors lacked the expected diffuse membranous CAIX expression and nuclear HIF2α staining typically observed in conventional ccRCC, consistent with incomplete activation of the canonical ccRCC hypoxia program **(Figure S14B)**. In agreement, WGS did not identify mutations affecting *VHL* or other chromosome 3p tumor suppressor genes, and tumor-derived cells largely retained expression of pVHL **(Figure S14C)**. These findings indicate that malignant transformation and metastatic potential can arise despite retention of the remaining *VHL* allele, suggesting that chromosome 3p loss together with recurrent adaptive aneuploidies can drive substantial tumor evolution prior to complete *VHL* inactivation.

Multiplex FISH revealed subtle karyotypic alterations during *in vivo* evolution. All tumors harbored partial chromosome 4p loss caused by non-reciprocal translocation with chromosome 22 and a translocation between chromosomes 10 and 13 **(Figure S14D)**. Additional chromosome-level alterations were detected in individual tumors (e.g., chromosome 15 loss in tumor 1, a non-reciprocal translocation between chromosomes 7 and 11 in tumor 2) **(Figure S14D)**. Thus, tumor progression was accompanied by continued karyotypic diversification *in vivo*, indicative of ongoing chromosomal instability.

Consistent with the metabolic adaptations observed *in vitro*, tumor-derived cells exhibited elevated levels of p62, indicative of persistent activation of the mTORC1 pathway **(Figure S14E)**. Moreover, tumor cells isolated from lung metastases displayed greater maximal respiratory capacity compared to the parental clone **(Figure 7E)**, demonstrating that enhanced OXPHOS is progressively selected during metastatic evolution, consistent with recent isotope-tracing studies demonstrating increased mitochondrial metabolism in human ccRCC metastases^59^.

Overall, these findings support a model in which chromosome 3p loss initiates a selective bottleneck characterized by p53 activation, oxidative stress, cell cycle arrest, and senescence **(Figure 7F)**. Rare cells that overcome this barrier undergo clonal expansion and continued genomic evolution, repeatedly acquiring chromosome 5q gain and 14q loss. This stepwise evolutionary process culminates in metabolic reprogramming, tumor formation, and elevated metastatic potential, thereby recapitulating the evolutionary trajectory of ccRCC from a single initiating chromosomal event. Thus, we establish a genetically defined platform for studying early ccRCC evolution and provide experimental evidence that stepwise chromosomal alterations can drive tumorigenesis.

## DISCUSSION

Tumorigenesis usually proceeds through iterative cycles of mutation and selection^78^. Here we describe a model in which a single catastrophic chromosomal event can reconfigure evolutionary potential. By inducing a targeted DSB on chromosome 3p, we show that 3p loss initiates clonal evolution in a punctuated manner. Rather than acting as a classical oncogenic driver that immediately enhances fitness, 3p loss imposes a profound proliferative burden while simultaneously reshaping the fitness landscape and creating new evolutionary trajectories. Cells acquiring 3p loss experience strong negative selection, thereby creating an evolutionary bottleneck in which most cells are eliminated or undergo permanent arrest. Only rare lineages that acquire compensatory alterations persist. Thus, tumor initiation reflects evolutionary escape from a growth-suppressive bottleneck rather than the immediate oncogenic effects of chromosome 3p loss itself.

Once this barrier is overcome, evolution proceeds through recurrent chromosomal alterations that mirror the genomic landscape of human ccRCC^16,79^. Independent clones repeatedly acquire chromosome 5q gain and 14q loss, the two most common secondary alterations in patient tumors, demonstrating that these events are not merely stochastic consequences of chromosomal instability but rather adaptive solutions to the fitness constraints imposed by 3p loss. Early ccRCC evolution thus resembles somatic punctuated evolution: chromosome 3p loss initially reduces cellular fitness, creating strong selective pressure for additional alterations that alleviate this burden and increase malignant potential. The repeated emergence of chromosome 14q loss across genetically distinct clones is particularly notable, providing direct experimental evidence that this alteration is a deterministic consequence of selection rather than a passenger event. Although *HIF1A* haploinsufficiency may partially contribute to this process, the selective advantage conferred by chromosome 14q loss remains incompletely understood and warrants further investigation.

On an evolutionary timescale, chromosome 3p loss and 5q gain are thought to arise simultaneously or in close temporal proximity during early ccRCC development, yet how these alterations originate and become coupled has remained unresolved^75^. In this study, we show that a single DSB on chromosome 3p is sufficient to generate both major classes of 3p alterations observed in ccRCC **(Figure 1)**. The simplest outcome is the loss of an acentric chromosome arm, producing the arm-level deletions commonly observed in patient tumors. Alternatively, replication of the broken chromosome followed by sister chromatid fusion generates a dicentric chromosome whose bridge fragmentation produces localized SVs confined to the original breakpoint region – a pattern we term breakpoint-confined chromothripsis **(Figures 1, 5A-B)**. This mechanism is consistent with recent studies demonstrating that persistent chromatin bridges can undergo localized fragmentation and repair, generating punctuated genome restructuring and sequence-level signatures such as tandem short template (TST) jumps^27,28^. Notably, while chromosome 3p fragmentation generated breakpoint-confined chromothripsis in our system, highly complex rearrangements characteristic of canonical chromothripsis were rarely observed. One possible explanation is that extensive chromosome fragmentation is poorly tolerated in non-transformed renal epithelial cells, resulting in strong negative selection. Additional breakage-fusion-bridge cycles can subsequently generate a derivative chromosome harboring concurrent 3p loss and 5q gain **(Figure 5C-D)**. Thus, both simple deletions and complex rearrangements characteristic of early ccRCC can arise from a single initiating lesion, providing a unified mechanistic explanation for the major classes of chromosome 3p alterations observed in human tumors **(Figure 5E)**.

Coincident with chromosome 14q loss, late-passage clones underwent metabolic reprogramming characteristic of ccRCC. Chromosome 3p loss initially imposed oxidative and metabolic stress, and surviving clones adapted through metabolic rewiring marked by enhanced mitochondrial respiration, mTORC1 activation, and upregulation of *SQSTM1*/p62. Interestingly, upregulation of chromosome 5q-encoded *SQSTM1*/p62 occurred independently of chromosome 5q gain, and its loss robustly impaired anchorage-independent growth, identifying p62 as a functional requirement of adaptation to chromosome 3p loss rather than a secondary consequence of chromosomal evolution. These observations suggest that activation of the p62–mTOR axis represents a general adaptive response to chromosome 3p loss. Together, these adaptations appear to buffer the fitness costs imposed by 3p loss and may represent therapeutically exploitable vulnerabilities that emerge during tumor evolution.

The evolutionary trajectory described above ultimately culminates in malignant transformation *in vivo*. Late-passage clones with 3p loss formed xenograft tumors and exhibited spontaneous metastatic dissemination to the lungs, demonstrating acquisition of malignant and metastatic phenotypes within a genetically defined system. Unexpectedly, transformation occurred despite retention of the remaining *VHL* allele and without the robust HIF activation that characterizes canonical ccRCC. While complete *VHL* inactivation likely remains necessary for the development of *bona fide* ccRCC, our findings demonstrate that substantial evolutionary progression can occur downstream of chromosome 3p loss prior to complete *VHL* loss. More broadly, our results suggest a generalizable principle by which deleterious initiating chromosomal alterations can reshape the fitness landscape and drive deterministic tumor evolution. By capturing the transition from a single chromosome break to metastatic disease, the isogenic models described here provide a tractable platform for identifying the genetic dependencies and therapeutic vulnerabilities that emerge during the earliest stages of cancer evolution.

## ACKNOWLEDGEMENTS

We thank Samra Turajlic for helpful discussions and Elizabeth Maurais, Kidist Woldehawariat, and Bing-Huan Wu for technical assistance. We acknowledge the Children’s Research Institute (CRI) Moody Flow Cytometry Core, CRI Metabolomics Facility (CPRIT RP240494), UT Southwestern Flow Cytometry Core, and the UT Southwestern Tissue Management Shared Resource of the Harold C. Simmons Comprehensive Cancer Center (NIH P30CA142543) for assistance and/or access to equipment and resources. We also thank Carlos Arteaga for shared use of equipment. This work was supported by the U.S. Department of Defense (W81XWH2210764 to PL), U.S. National Institutes of Health (R35GM146610 and R01CA289435 to PL; P50CA196516 to JB; R01CA294636 to QZ; R01CA258629 to SM), Cancer Prevention and Research Institute of Texas (RR180050 to PL), UT Southwestern Kidney Cancer Program, UT Southwestern Circle of Friends (to PL), Welch Foundation (I-2071-20240404 to PL), and European Molecular Biology Laboratory core funding (to IC-C). We acknowledge the computational resources provided by the EMBL European Bioinformatics Institute.

## AUTHOR CONTRIBUTIONS

Conception: RD, IC-C, PL Design: RD, IVB, IC-C, PL

Performed experiments: RD, CL, Y-FL, AK (Ko), AKM, JZ, QH, JLE, JM

Data analysis: RD, IVB, AK (Kumar), Y-FL, AK (Ko), AKM, EA, QH, JLE, JEV-I, AGR, PK, IC-C

Supervision: RM, JB, SM, SZ, PK, QZ, IC-C, PL

Funding: JB, SM, QZ, IC-C, PL

Writing: RD, IVB, IC-C, PL with input from all authors

## DATA AVAILABILITY STATEMENT

All data reported in this study are available within the main text and supplementary materials. Whole-genome sequencing (WGS) data from TCGA are available under controlled data access through dbGaP under accession code dbGaP:phs000178. WGS data from Senkin et al. are available through the European Genome-Phenome Archive (EGA) under accession number EGAS00001003542. WGS data generated in this study have been deposited in the European Nucleotide Archive (ENA) at EMBL-EBI under accession number PRJEB114979. Bulk RNA sequencing and single-cell RNA sequencing data generated in this study have been deposited in the NCBI Gene Expression Omnibus (GEO) under accession numbers GSE336576 and GSE336056, respectively.

## METHODS

### Cell lines and reagents

Cells were maintained at 37°C in a humidified atmosphere containing 5% CO_2_. RPTECs were obtained from ATCC (CRL-4031) and cultured in renal epithelial cell growth basal medium (Lonza) supplemented with 5% tetracycline-free fetal bovine serum (Omega Scientific), 100 U mL^-1^ penicillin-streptomycin, 10 ng mL^-1^ human recombinant epidermal growth factor, 1x insulin-transferrin-selenium solution, 1 μg mL^-1^ hydrocortisone, 10 μM epinephrine, 50 ng mL^-1^ triiodo-L-thyronine, 30 μg mL^-1^ gentamicin, and 15 ng mL^-1^ amphotericin B. RPTECs were further modified to express CDK4 and a short hairpin RNA targeting *TP53*, as indicated in the text. 786-O, HEK293T, and 293GP cells were cultured in Dulbecco’s modified Eagle medium (DMEM; Gibco) supplemented with 10% tetracycline-free fetal bovine serum (Omega Scientific) and 100 U mL^-1^ penicillin-streptomycin. All cell lines were authenticated by karyotyping and routinely tested for mycoplasma contamination using the Universal Mycoplasma Detection Kit (ATCC), with all results confirmed to be negative.

To induce chromosome mis-segregation and micronuclei formation by silencing of the mitotic spindle assembly checkpoint, cells were treated with 500 nM Mps1 inhibitor (NMS-P715; Cayman Chemical). For ionizing radiation experiments, cells were exposed to the indicated doses of γ-irradiation using a Mark I^137Cs^ irradiator (JL Shepherd) and fixed 1 hr post-irradiation for immunofluorescence analysis. DMOG (MilliporeSigma) was dissolved in DMSO and used at the indicated concentrations. Geneticin (G418 sulfate; InvivoGen), hygromycin (Hygromycin B Gold; InvivoGen), puromycin (Puromycin dihydrochloride; Sigma-Aldrich) and blasticidin (Blasticidine S hydrochloride; Sigma-Aldrich) were used at 100, 300, 2.5 and 10 μg mL^-1^, respectively. For colony staining, MTT (5 mg mL^-1^ stock in PBS) was diluted to a final concentration of 1 mg mL^-1^. To suppress non-homologous end joining (NHEJ), the DNA-PKcs inhibitor AZD7648 (MedChemExpress) was used at a concentration of 1 μM. For cell cycle arrest, cells were treated with nocodazole (100 ng mL^-1^; MilliporeSigma) or Colcemid (100 ng mL^-1^; KaryoMAX, Thermo Fisher Scientific), prepared in DMSO. dTAGv1 (Tocris Bioscience) was used at 0.5 μM to induce degradation of FKBP12^F36V^ degron-tagged proteins.

### Generation of engineered cell lines

Cas9 (TrueCut Cas9 protein v2; Invitrogen or Alt-R S.p. dCas9 Protein V3; IDT) ribonucleoprotein (RNP) complexes were delivered using Lipofectamine CRISPRMAX Cas9 Transfection Reagent (Invitrogen) according to the manufacturer’s instructions. Target sequences for guide RNAs were designed using CRISPick (Broad Institute) or CRISPOR and synthesized by GenScript. sgRNA sequences used in this study were: sg3p-1, ATTACCTTGGGCCCAACATA; sg3p-2, ACTTAGGAATTAACCGCGAC; sgNTC, GCTTAGTTACGCGTGGACGA; sgYq, AACACTTCTCTAGCACGATT; sgVHL, CCCGTATGGCTCAACTTCGA; sg*RB1*, TGAACTACTTACGAACTGCT; sg*SQSTM1*-1, GGCTTCCAGGCGCACTACCG; sg*SQSTM1*-2, TGGCTCCGGAAGGTGAAACA; and sg*SQSTM1*-3, ATATCGATGTGGAGCACGGA.

Retroviral particles were produced in HEK293GP cells by co-transfection of transfer plasmid with pVSV-G (139479, Addgene) using X-tremeGENE 9 (Millipore-Sigma). Viral supernatants were collected 48 and 72 hr post-transfection, filtered (0.45 μm), and used to infect target cells supplemented with 8 μg mL^−1^ polybrene (Millipore-Sigma) for around 24 hr. Lentiviral particles were produced in HEK293T cells by co-transfection of transfer plasmid, psPAX2 (12260, Addgene), and pMD2.G (12259, Addgene) using X-tremeGENE 9. Viral supernatants were collected at 48 and 72 hr, filtered, and used to infect target cells supplemented with 8 μg mL^−1^ polybrene for ∼24 hr. For transient expression, cells were transfected using X-tremeGENE 9. KO clones were expanded and verified by both Sanger sequencing and immunoblotting.

### Fluorescence-activated cell sorting (FACS)

Cells were harvested by trypsinization, washed once with phosphate-buffered saline (PBS), and resuspended at 2 × 10^6^ cells mL^-1^ in sorting buffer (PBS supplemented with 2% fetal bovine serum). Cell suspensions were filtered through mesh-capped polypropylene tubes immediately prior to sorting. Sorting was performed using a FACSAria II cell sorter (BD Biosciences) equipped with a 100 μm nozzle and controlled by FACSDiva software (v8.0.2). Debris was excluded by forward- and side-scatter gating, and doublets were removed using FSC-A versus FSC-W gating. For dTAG-Flow experiments, GFP-positive gates were established using untreated GFP-degron-expressing cells and validated against non-fluorescent parental controls. For clonal isolation, single cells were sorted directly into 96-well plates containing pre-warmed complete growth medium. Data were analyzed using FlowJo (v10).

### CRISPR-Cas9-mediated chromosome breakage

RNP complexes were assembled *in vitro* immediately prior to transfection. sgRNAs were combined with Cas9 at a molar ratio of 3:1 (sgRNA:Cas9) and incubated at room temperature for 8 min to allow for complete RNP assembly using Lipofectamine CRISPRMAX Cas9 transfection reagent according to the manufacturer’s instructions. Either DMSO or 1 μM AZD7648 was supplemented into the culture medium at the time of transfection. At 24 hr post-transfection, the culture medium was replaced with complete medium containing AZD7648 to sustain inhibitor activity throughout the active editing window. At 48 hr post-transfection, the medium was replaced with complete medium without AZD7648. At 72 hr post-transfection, cells were processed for cytogenetic analysis. Micronuclei frequency and chromosomal alterations were assessed by FISH using chromosome paint and locus-specific FISH probes from interphase cells and metaphase spreads, respectively, enabling quantification of Cas9-induced alterations of the targeted chromosome.

For chromosome bridge analysis, cells were processed to distinguish mitotic and interphase bridges. For mitotic bridge analysis, 8 hr prior to the 72 hr time point, cells were treated with nocodazole for 6 hr to enrich for mitotic cells, followed by three washes with fresh media (5 min each) to remove the drug. Cells were released from nocodazole for 105-120 min to allow progression into anaphase or telophase, fixed with formaldehyde, and processed for FISH. For interphase bridge analysis, cells were fixed at 72 hr time point without nocodazole treatment to preserve native interphase structures, followed by the same fixation and FISH procedure.

### Metaphase chromosome spread preparation

Metaphase spreads were prepared by treating cells with 100 ng mL^-1^ colcemid (KaryoMAX, Thermo Fisher) for 4 hr to enrich for mitotic cells. Cells were then trypsinized, pelleted by centrifugation, and the supernatant was discarded before gently resuspending the pellet in residual medium by flicking. For hypotonic treatment, 5 mL of pre-warmed 0.075 M KCl was added dropwise while slowly vortexing to prevent cell lysis. Cells were incubated at 37°C for 6 min to allow swelling. Subsequently, 1 mL of cold Carnoy fixative (3:1 methanol:acetic acid) was added, mixed gently by inversion, and the cells were centrifuged at 180 x g for 5 min and the supernatant removed. The fixation step was repeated by resuspending the pellet in 5 mL of cold Carnoy fixative, followed by gentle mixing, centrifugation, and removal of the supernatant.

The final cell pellet was resuspended in approximately 200 μL of cold Carnoy fixative (volume adjusted as needed). Fixed cell suspensions were stored at -20°C for long-term use. For slide preparation, 10 μL of the suspension was dropped onto a slide and allowed to air-dry and metaphase spreads were examined under a microscope.

### DNA fluorescence *in situ* hybridization (FISH)

DNA FISH probes (MetaSystems) were applied to metaphase chromosome spreads and sealed with a coverslip using rubber cement. Slides were co-denatured at 75°C for 2 min and hybridized overnight at 37°C in a humidified chamber. After hybridization, slides were washed in 0.4x SSC at 72°C for 2 min, followed by 2x SSC with 0.05% Tween-20 at room temperature for 30 seconds, then counterstained with DAPI and mounted in ProLong Gold. For multiplex FISH, metaphase spreads were hybridized with 24XCyte (MetaSystems) probes. For locus-specific DNA FISH targeting chromosome 3p, the RP11-447A21 BAC clone was obtained from BACPAC Resources, purified using a NucleoBond Xtra BAC kit (Macherey-Nagel), and labeled by nick translation using the DNA Labeling System 2.0 (Enzo Life Sciences). FISH was then performed using these probes as described above. For telomere FISH, slides were denatured at 80°C, and the G-rich telomeric strand was detected using a TelG telomere PNA probe (PNA Bio),

### Oligopaint DNA FISH

Oligopaint probes were prepared from pooled libraries by PCR amplification using Phusion Hot Start Master Mix and primers containing a T7 promoter sequence. Amplified products were purified using DNA Clean & Concentrator columns (Zymo Research) and subjected to in vitro transcription using the HiScribe T7 system (New England Biolabs). RNA products were reverse transcribed using fluorophore-labeled primers and Maxima H Minus reverse transcriptase (Thermo Fisher Scientific), followed by RNA hydrolysis and purification. Final DNA probes were quantified prior to use.

For Oligopaint FISH, we used a modified protocol described by Beliveau et al. ^80^. Cells were permeabilized in freshly prepared buffer (1x PBS, 0.5% Triton X-100) for 10 min at room temperature, washed in PBT, and treated with 0.1 N HCl for 5 min. Samples were rinsed in 2x SSCT and equilibrated in 2x SSCT with 50% formamide, followed by incubation at 60°C for 20 min. Oligopaint probes (∼1.6 μM) were prepared in hybridization buffer, applied to samples, denatured at 78°C for 3 min, and hybridized at 37°C for 16-24 hr. After primary hybridization, samples were washed in 2x SSCT at room temperature and 60°C, followed by additional washes in 2x SSCT and 0.2x SSCT. Secondary hybridization was performed using labeled probes in a 25 μL reaction for 30-120 min at room temperature. Post-hybridization washes were carried out as above, after which samples were stained with DAPI, rinsed, and mounted. Imaging was performed using the Metafer Scanning and Imaging Platform (MetaSystems), as described below.

### Fluorescence microscopy

Immunofluorescence and FISH imaging were performed on a DeltaVision Ultra microscope system (Cytiva/GE Healthcare) equipped with a 4.2 MP sCMOS detector. Interphase cells, micronuclei, and metaphase spreads were imaged using a 100x oil objective (UPlanSApo, 1.4 NA) or a 60x oil objective (PlanApo N, 1.42 NA). Z-stacks were acquired at 0.2 μm intervals (typically 11-15 sections) and deconvolved maximum intensity projections were generated using softWoRx (v7.2.1, Cytiva). Quantitative image analysis was performed using Fiji (v2.1.0/1.53c). Metaphase DNA FISH images were acquired using the Metafer Scanning and Imaging Platform (Metafer 4, v3.13.6, MetaSystems). Slides were initially scanned for metaphases using M-search with a 10x objective (ZEISS Plan-Apochromat 10x/0.45), followed by automated image acquisition with Auto-cap using a 63x oil objective (ZEISS Plan-Apochromat 63x/1.40). Images were analyzed using Isis (MetaSystems) and Fiji.

### Immunoblotting

Whole-cell lysates were prepared in Laemmli SDS sample buffer and heated at 95°C for 10 min. Proteins were separated by SDS-PAGE and transferred onto 0.2 μm PVDF membranes, which were subsequently blocked in 5% milk prepared in PBST (PBS containing 0.1% Tween-20). Membranes were incubated with primary antibodies diluted in PBST, including anti-pVHL (1:1,000; Cell Signaling), anti-α-tubulin (1:5,000; Cell Signaling), anti-β-actin (1:5,000; Cell Signaling), anti-HIF1α (1:1,000; Cell Signaling), anti-HIF2α (1:1,000; Cell Signaling), anti-p53 (1:1,000; Santa Cruz), anti-phospho-p53 (S15) (1:1,000; Cell Signaling), anti-Rb (1:1,000; Cell Signaling), anti-SOD3 (1:1,000; Cell Signaling), anti-HA (1:1,000; Novus), anti-SETD2 (1:1,000; Cell Signaling), anti-BAP1 (1:1,000; Cell Signaling), anti-PBRM1 (1:1,000; Bethyl), anti-p62 (1:1,000; Santa Cruz), anti-phospho-4E-BP1 (S65) (1:1,000; Cell Signaling), and anti-phospho-S6 K1 (T389) (1:1,000; Cell Signaling). After primary antibody incubation, membranes were briefly rinsed three times with PBST for 3 x 15 min. After washing, membranes were probed with HRP-conjugated secondary antibodies (goat anti-rabbit or donkey anti-mouse, Invitrogen) diluted 1:4,000 in 5% milk for 1 hr at room temperature. Signal detection was carried out using SuperSignal West Pico Plus chemiluminescent substrate (Thermo Fisher Scientific), and images were acquired with a ChemiDoc MP imaging system (Bio-Rad).

### Cell proliferation and transformation assays

Anchorage-independent growth was assessed using a soft agar colony formation assay. Briefly, a base layer of 0.8% agar in complete growth medium was prepared in 6-well plates and allowed to solidify at room temperature. Cells were then suspended in 0.4% agar in complete medium and seeded on top of the base layer at a density of 5000 cells per well. After solidification, culture medium was added to prevent drying and replenished every 3-4 days.

Cells were incubated at 37°C in a humidified atmosphere with 5% CO_2_ for 4 weeks to allow colony formation. Colonies were stained by adding 500 μL of MTT solution (5 mg mL^-1^ in PBS) to each well and incubated at 37°C for 2-4 hr. After incubation, the staining solution was removed, and 500 μL of PBS added to each well. Plates were imaged and quantified using a GelCount colony counter (Oxford Optronix). Colonies larger than ∼70 μm in diameter were counted.

Cell proliferation was assessed by seeding 2 x 10^5^ cells per well in triplicate in 6-well plates and counting cells at 3-day intervals. For clonogenic survival assays, 100-200 cells were plated per well in triplicate in 6-well plates and cultured for 10 days. Colonies were then fixed with 100% ethanol and stained with 0.5% crystal violet in 70% ethanol. Colonies were either manually counted or scanned using a LI-COR imaging system (Odyssey). For LI-COR-based analysis, plates were imaged under standardized exposure settings, and colony area and integrated intensity per well were quantified using LI-COR image analysis software.

### Mitochondrial respiration measurements

Oxygen consumption rates (OCR) were measured using a Seahorse XFe96 Analyzer (Agilent Technologies). Cells were seeded at 1 x 10^4^ cells per well in Seahorse XF96 cell culture microplates 24 hr prior to the assay to allow attachment and recovery. On the day of the assay, culture medium was replaced with Seahorse XF assay medium supplemented as appropriate with glucose, pyruvate, and glutamine, and cells were equilibrated in a non-CO_2_ incubator at 37°C for 30-45 min. OCR measurements were obtained under baseline conditions followed by sequential injections of mitochondrial inhibitors: 2 μM oligomycin A, 1 μM carbonyl cyanide 3-chlorophenylhydrazone, and 2 μM antimycin A. Measurements were recorded at multiple time points following each injection according to the manufacturer’s instructions. Cell numbers per well were determined using a Celigo Image Cytometer (Revvity), and OCR values were normalized accordingly. Data was analyzed using Seahorse Wave software.

### Whole-genome DNA sequencing analysis

WGS data from Senkin et al.^37^ were downloaded from the EGA with accession ID EGAS00001003542. TCGA WGS data were downloaded from the Genomics Data Commons Data Portal (https://portal.gdc.cancer.gov). For TCGA, we restricted our analysis to cases with WGS data comprising paired-end reads exceeding 100 base pairs in length. For cases with multiple primary tumour BAM files available, we selected the BAM file with the most recent date of creation. Raw sequencing reads from both cohorts were mapped to the hg38 build of the human reference genome using BWA-MEM version 0.7.17-r1188. Aligned reads in BAM format were processed following the Genome Analysis Toolkit (GATK, version 4.1.8.0) Best Practices workflow to remove duplicates and recalibrate base quality scores^81,82^. Somatic SVs were called with GRIDSS2^83^ (v2.12.0, https://github.com/PapenfussLab/gridss), annotated with RepeatMasker (v4.1.2-p1, http://www.repeatmasker.org) and kraken2^84^ (v2.1.2), filtered with GRIPSS (v1.9), and clustered and visualized with LINX (v1.15)^85^. B-allele frequency (BAF) values for heterozygous SNP sites were computed using AMBER (v3.5) and read depth ratios were calculated with COBALT (v1.11). B-allele frequency information, read-depth ratios, SV breakpoints and allele frequencies of SNVs were integrated to estimate the purity, ploidy and copy-number profile of each tumor using PURPLE (v2.54)^85^. Both minor and total copy-number values computed using PURPLE were rounded to integer values. Finally, somatic and germline SNVs, multi-nucleotide variants and indels were called and filtered using SAGE (v2.8). Quality control was performed using AMBER and PURPLE. Samples were discarded when: (1) FAIL_CONTAMINATION: measured tumor contamination in homozygous sites from the normal sample was >10%; (2) FAIL_NO_TUMOR: no evidence of tumor was found in the sample – in these cases, samples were kept if at least one of the following criteria were met: (i) the tumor had one or more HOTSPOT SV or point mutations, (ii) the number of somatic SNVs was > 1000, (iii) the number of somatic SVs was > 1000 (excluding single breakends), or (iv) the tumor sample had 3000 BAF points in germline diploid regions with a tumor ratio < 0.8 OR > 1.2 (evidence of some level of aneuploidy). SAGE, GRIPSS, AMBER, COBALT, PURPLE and LINX are developed by the Hartwig Medical Foundation and code, tools and documentation are freely available at: https://github.com/hartwigmedical/hmftools.

### Detection of breakpoint-confined chromothripsis

Visual inspection of rearrangement profiles recurrently detected in chromosome 3p in ccRCC tumors, which we term ‘≈-confined chromothripsis’, were characterized by the presence of: (1) cluster of SVs, often including translocations, confined to a small chromosomal region relative to the chromosome size; (2) copy number oscillations in the regions encompassed by the SV cluster; and (3) terminal loss. To classify the rearrangement mechanisms underpinning 3p loss in a data-driven manner, we used a stepwise approach considering the presence of SV clusters, copy-number oscillations and terminal loss. First, tumors were classified based on 3p loss status into no loss, full chromosome 3 loss and 3p loss. Subsequently, tumors with 3p loss were classified into canonical chromothripsis using ShatterSeek v1.1^16,29^, breakpoint-confined chromothripsis, and ‘other 3p loss’. Next, we applied ShatterSeek v1.1 to identify clusters of SVs, requiring at least six SVs of which at least four had to be interleaved. In addition, we considered chromosomes linked by at least three interchromosomal SVs to be involved in an SV cluster. Tumors were considered to have confined chromothripsis on 3p if at least one of the following sets of criteria was satisfied: (1) SV cluster smaller than 14.7 Mbp and terminal loss of at least 1 Mbp and mapping less than 5Mbp away from the SV cluster; or (2) presence of at least one translocation linking chromosome 3, with or without copy-number oscillations, with an SV cluster on another chromosome and terminal loss on 3p of at least 1 Mbp and mapping less than 5Mbp away from the most upstream translocation breakpoint. To determine the SV cluster size threshold, we applied an expectation–maximization approach. Specifically, we applied the Mclust function from the R package mclust to SV clusters mapping to chromosome 3p across all cancer types from the TCGA cohort analyzed in this study. This analysis revealed that 85% of SV clusters detected mapping to chromosome 3p in ccRCC tumors spanned less than 14.7 Mpb.

Because canonical chromothripsis events involving a small number of SVs, as is often the case in ccRCC tumors^16,29^, might not meet the criteria for chromothripsis detection, we performed manual inspection of rearrangement and copy-number profiles for all cases^29^. Using these criteria, we achieved a precision of 97% and recall of 78% for the classification of confined chromothripsis when manually inspecting the SV clusters detected in 949 ccRCC tumours from the Senkin and TCGA cohorts. Rearrangement and SCNA profiles were visualized using ReConPlot^86^ (https://github.com/cortes-ciriano-lab/ReConPlot).

### Single-cell RNA sequencing and analysis

RPTECs were transfected with sgNTC, sgYq, or sg3p in duplicates and harvested on days 5 and 10. To generate single-cell suspensions, cells were trypsinized and were collected by centrifugation at 300 x g for 5 min at 4°C. Cell pellets were washed twice with PBS and resuspended in PBS supplemented with 0.04% BSA. The cell suspension was passed through a 40 μm cell strainer to remove aggregates. Cell viability was assessed using trypan blue exclusion for downstream applications. Single-cell suspensions were immediately used for library preparation following the 10x Genomics protocol.

#### Multiplexing of samples for single-cell profiling

To enable simultaneous transcriptomic profiling and sample multiplexing, cells from distinct conditions were labeled with unique oligonucleotide-conjugated hashtag antibodies according to the CITE-seq STAR protocol^87^. Briefly, cells were resuspended at 1 x 10^6^ cells in 100 μL staining buffer (2% BSA, 0.02% Tween-20 in PBS), followed by Fc receptor blocking using Fc Blocking Reagent (FcX, BioLegend) for 20 min at 4°C. Cells were then incubated with TotalSeq-A hashtag antibodies (BioLegend) for 30 min at 4°C. After staining, cells were washed extensively with CITE-seq Wash Buffers A, B, and C, including an additional final wash in Buffer C to ensure complete removal of unbound antibodies^87^. Cells were subsequently resuspended in PBS at 1,800 cells/μL, pooled and immediately processed for GEM generation using the 10x Genomics Chromium Controller, targeting a recovery of 10,000 cells. The following BioLegend TotalSeq-A hashtag antibodies were used: HTO-1 (394601), HTO-2 (394603), HTO-3 (394605), HTO-4 (394607), HTO-5 (394609), and HTO-6 (394611).

#### scRNA-seq library construction and sequencing

Reverse transcription and transcriptome library preparation were performed using the 10x Genomics Chromium Single Cell 3′ Reagent Kit v3.1 (CG000205) according to the manufacturer’s instructions. Hashtag oligonucleotides (HTO) library fractions were isolated following initial cDNA amplification (Step 2: Post-GEM RT Cleanup & cDNA Amplification) and processed separately from the whole-transcriptome libraries. In brief, HTO libraries were generated using a standard CITE-seq hashing workflow (https://citeseq.files.wordpress.com/2019/02/cite-seq_and_hashing_protocol_190213.pdf). Following amplification, libraries were indexed using Illumina-compatible adapters. The quality and quantity of transcriptome and HTO libraries were assessed using Qubit fluorometry and Agilent TapeStation prior to sequencing. Libraries were pooled and sequenced on an Illumina NovaSeq X Plus platform using paired-end 150 bp reads (PE150).

#### Data processing and analysis

FASTQ files were generated using Cell Ranger mkfastq (v7.2.0, 10x Genomics). Gene expression reads were aligned to the GRCh38-2020-A reference genome and quantified using Cell Ranger count (v7.2.0). HTO reads were quantified using CITE-seq-Count, and sample demultiplexing was performed using the HTODemux function in Seurat (v4.0). Quality control filtering excluded cells with high mitochondrial gene content, low gene counts, or ambiguous hashtag assignments. Following filtering, data were normalized and scaled prior to dimensionality reduction and clustering. Where applicable, batch effects were corrected using Harmony or Seurat SCTransform-based integration. Volcano plots were generated using ggplot2, and genes were labelled using ggrepel. For GSEA, genes were ranked by average log2 fold change (avg_log2FC), and GSEA was performed using the fgsea R package with the multilevel algorithm and MSigDB Hallmark and SenMayo gene sets. Enrichment plots were generated using fgsea, and pathways with an FDR-adjusted *P* value < 0.05 were considered significantly enriched.

### Bulk RNA sequencing and analysis

RNA-seq libraries were prepared and sequenced on an Illumina NovaSeq platform, generating paired-end 150 bp reads. Raw sequencing reads were processed using Trim Galore (v0.6.10), which incorporates Cutadapt (v2.5). Adapter trimming and quality filtering included removal of 10 bp from the 5′ end and 1 bp from the 3′ end of each read. Reads shorter than 25 bp after trimming were discarded. Trimming was performed in paired-end mode using six threads. Trimmed reads were aligned to the human reference genome (GRCh38, GENCODE release 37) using STAR (v2.7.2b) in two-pass mode to improve splice junction detection. Alignment was performed using eight threads with relaxed filtering to retain partially aligned reads. Overall alignment rates ranged from 83% to 90% across samples. Aligned reads were processed using samtools (v1.22.1). Properly paired reads with MAPQ ≥ 255 were retained and sorted by read name for downstream quantification.

Gene-level counts were generated using featureCounts (Subread v2.0.6) by assigning reads to exonic regions based on GENCODE release 37 annotations. Paired-end counting was enabled with a minimum mapping quality threshold of 20. Data were analyzed in an unstranded mode. Downstream analyses were performed in R (v4.3.3). Ensembl gene identifiers were converted to gene symbols, and duplicate genes were collapsed by summing counts. Genes with low expression (total counts < 30 across all samples) were excluded. Differential expression analysis was performed using DESeq2 (v1.42.1), modeling counts using a negative binomial distribution with the design formula ∼ clone_L. P values were adjusted for multiple testing using the Benjamini–Hochberg method. Genes with adjusted P < 0.05 and log2 fold change > 1 were considered significantly differentially expressed.

For GSEA, genes were ranked by the Wald statistic from DESeq2. Duplicate gene symbols were collapsed by averaging. GSEA was performed using the fgsea (v1.28.0) with the multilevel algorithm and MSigDB Hallmark gene sets using msigdbr (v25.1.1). Pathways with adjusted P < 0.05 were considered significant. The top 10 positively and negatively enriched pathways were selected based on normalized enrichment score (NES). Dot plots were generated using ggplot2, with dot size representing gene set size and color indicating −log10(adjusted P value). Statistical analyses were performed in R (v4.3.3), and unless otherwise stated, adjusted P < 0.05 was considered statistically significant. For gene expression Z-score analysis, DESeq2-normalized counts were log2-transformed [log2(normalized counts + 1)], and gene-wise Z-scores were calculated in R.

### Xenograft tumor studies

All animal experiments were performed in accordance with US National Institutes of Health (NIH) guidelines and approved by the Institutional Animal Care and Use Committee (IACUC) at UT Southwestern Medical Center (protocol no. 2019-102794). Mice were maintained under specific pathogen-free conditions in the UT Southwestern Animal Resource Center. RPTECs were cultured under standard conditions and harvested during logarithmic growth. Cells were washed with sterile PBS and resuspended on ice in a 1:1 mixture of PBS and Matrigel (354234, Corning). A total 2 x 10^6^ cells in 100 μL were injected subcutaneously into each dorsal flank (bilateral injection) of 6-8-week-old male NOD scid gamma (NSG) mice (Jackson Laboratory). Tumor dimensions were measured serially using digital calipers, and tumor volume was calculated as (length x width^2^)/2. Mice were euthanized upon reaching predefined ethical or tumor-size endpoints, and tumors were collected immediately for downstream analyses.

### Histology and immunohistochemistry (IHC)

IHC analyses were performed by the UT Southwestern Kidney Cancer Program Core Histology Facility on whole tumor tissue sections. Tissues were fixed in 10% neutral-buffered formalin, paraffin-embedded (FFPE), sectioned at 3-5 μm, and stained with hematoxylin and eosin (H&E) using standard protocols^88^. Antibodies used included PAX8, (clone MRQ-50, 363M-14, Cell Marque), HIF2α (clone A-5, SC-46691, Santa Cruz), and CA9 (PA1-16592, Invitrogen). IHC was performed using a Dako automated staining system (Agilent).

### Isolation of tumor-derived RPTECs

Primary tumors and lung metastases were harvested immediately following euthanasia and washed in ice-cold sterile PBS. All subsequent procedures were performed under sterile conditions. Tissues were mechanically minced into ∼1 mm^3^ fragments and digested in serum-free DMEM containing collagenase type IV (1 mg mL^-1^) at 37°C for 1-2 hr with gentle agitation. Digested tissue was pelleted by centrifugation (200 x g, 3 min), washed three times with PBS, and further dissociated in 0.25% trypsin-EDTA at 37°C for 5 min. Following centrifugation and an additional PBS wash, cells were resuspended in complete growth medium supplemented with 2x penicillin-streptomycin, filtered through a 100 μm EZFlow cell strainer, and plated in 10 cm culture dishes. Medium was replaced daily for the first 3 days to remove non-adherent cells and debris, and cultures were subsequently expanded under standard growth conditions. GFP-positive tumor-derived cells were isolated by FACS as described above.

### Quantification and statistical analysis

Statistical tests were performed as described in the figure legends using GraphPad Prism version 9.5.0. Sample sizes, statistical analyses, and significance values are reported in the figure legends, denoted in the figure panel, or described in the text. Definitions for n are described in the figure legends. All experiments were performed independently at least three times unless otherwise indicated. For statistical analyses, *P* > 0.05 was considered not significant (ns), and asterisks denote the following: \**P* ≤ 0.05; \*\**P* ≤ 0.01; \*\*\**P* ≤ 0.001; \*\*\*\**P* ≤ 0.0001. Error bars represent standard error mean (SEM) unless otherwise indicated.

## SUPPLEMENTARY FIGURE LEGENDS

**Figure S1. Additional genomic features of breakpoint-confined chromothripsis in ccRCC, related to Figure 1.**

**(A)** Mutation status of *VHL* was inferred from combined analysis of SNVs, SCNAs and SVs. Biallelic disruptions include SNVs with loss of heterozygosity or homozygous deletions. Monoallelic losses include single-hit SNVs, SV breakpoints mapping within genes, and hemizygous deletions of the full gene. Other SCNAs include partial hemizygous losses and gains.

**(B)** Frequency of biallelic *VHL* inactivation across TCGA cancer types.

**(C)** Frequency of chromosome 3 loss mechanisms across TCGA cancer types.

**(D)** Rearrangement profile showing breakpoint-confined chromothripsis involving chromosomes 3 and 6. Profiles display total somatic copy-number values and SV junctions. Total and minor copy-number values are shown in black and blue, respectively. DEL, deletion-like rearrangement; DUP, duplication-like rearrangement; h2hINV, head-to-head inversion; t2tINV, tail-to-tail inversion.

**Figure S2. Renal proximal tubule epithelial cells provide a genomically stable platform for engineering chromosome 3p alterations, related to Figure 2.**

**(A)** Cumulative population doublings of RPTECs with and without CDK4 expression.

**(B)** Clonogenic survival of RPTECs with and without CDK4 expression. Right, representative crystal violet-stained colonies.

**(C)** Multiplex FISH karyotype of RPTECs. A total of 90 metaphase spreads were analyzed.

**(D)** Frequency of micronuclei in RPTEC following treatment with an Mps1 inhibitor (Mps1i). Data were derived from analysis of 533 DMSO-treated and 1002 Mps1i-treated cells.

**(E)** Experimental workflow for Cas9-induced chromosome 3p breakage.

**(F)** Interphase FISH images showing chromosome 3-positive micronuclei 72 hr after Cas9-sgRNA RNP delivery with or without detectable chromosome 3-specific centromeric sequences (CEN3). Right, quantification of acentric or centromere-containing, chromosome 3-positive micronuclei. Scale bar, 5 µm.

**(G)** Interphase FISH images showing chromosome 3-positive micronuclei 72 hr after lentiviral Cas9-sgRNA expression. Scale bar, 5 µm.

**(H)** Quantification of micronuclei containing the indicated chromosomes as in (G). Data are mean ± SEM of *n* = 2 independent experiments from 881 EV cells and 949 sg3p-1 cells.

**(I)** Quantification of acentric or centromere-containing, chromosome 3-positive micronuclei. Data are from two independent experiments.

**(J)** Examples of chromosome 3-positive chromatin bridges following chromosome 3p targeting. Scale bar, 5 µm.

**Figure S3. p53 suppression promotes persistence of chromosome 3p alterations.**

**(A)** Oligopaint design spanning the proximal and distal regions of chromosome 3p and the 3q22 locus. Right, metaphase FISH image showing chromosome 3 labeling with Oligopaint probes. Scale bar, 10 µm.

**(B)** Metaphase FISH images showing chromosome 3p alterations detected by Oligopaint probes following Cas9-sgRNA RNP delivery. Scale bar, 10 µm.

**(C)** Immunoblot analysis of RPTECs expressing control or shp53 following irradiation.

**(D)** Clonogenic survival of RPTECs expressing shp53 following exposure to the indicated doses of ionizing radiation.

**(E)** Quantification of metaphase spreads from RPTECs expressing control or shp53 showing chromosome 3p alterations. Data are mean ± SEM from *n* = 3 experiments from 749 metaphase spreads from RPTECs and 252 metaphase spreads from RPTECs expressing shp53. Statistical analysis by two-way ANOVA test with multiple comparisons.

**Figure S4. Physiologically relevant ccRCC-associated conditions fail to select for chromosome 3p loss.**

**(A)** Immunoblot analysis showing HIF2α stabilization following growth under hypoxia (1% O_2_).

**(B)** Immunoblot analysis showing HIF2α stabilization following 6-hr treatment in the indicated concentrations of dimethyloxalylglycine (DMOG).

**(C)** Crystal violet-stained colonies following 14-day growth in the indicated concentrations of DMOG.

**(D)** Quantification of chromosome 3p loss following growth under normoxic or hypoxic conditions. Data were derived from 84-232 metaphase spreads per condition.

**(E)** Top, experimental workflow for assessing anchorage-independent growth following induction of chromosome 3p loss. Bottom, images of soft agar colony formation assays following chromosome 3p targeting. Scale bar, 1 mm.

**(F)** Quantification of anchorage-independent growth shown in (E).

**(G)** Quantification of chromosome 3p loss following growth under low-attachment conditions. Data are mean ± SEM from *n* = 2 experiments from 207-337 metaphase spreads analyzed per condition. Statistical analysis by two-way ANOVA test with multiple comparisons.

**(H)** Immunoblot analysis confirming generation of VHL knockout clones.

**(I)** Clonogenic survival assays of *VHL* WT and *VHL*^-/-^ cells. Right, quantification of colony formation.

**(J)** Quantification of chromosome 3p loss in *VHL* WT and *VHL*^-/-^ cells at the indicated time points. Data are mean ± SEM from *n* = 3 experiments from 39-273 metaphase spreads analyzed per condition or clone.

**(K)** Cumulative population doublings of *VHL* WT and *VHL*^-/-^ cells following chromosome 3p targeting.

**Figure S5. Single-cell transcriptomics reveal immediate transcriptional responses to chromosome 3p loss.**

**(A)** InferCNV analysis of cells following sgNTC, sgYq, or sg3p targeting at days 5 and 10 after Cas9-sgRNA RNP delivery.

**(B)** Frequency of cells harboring chromosome 3p loss identified by InferCNV at the indicated timepoints.

**(C)** Relative expression of chromosome 3 genes in chromosome 3p loss cells. Red arrow indicates the Cas9 cut site.

**(D-E)** Volcano plots showing differentially expressed genes in chromosome 3p loss cells relative to cells without chromosome 3p loss day 5 (D) and day 10 (E). Differentially expressed chromosome 3p genes are indicated.

**Figure S6. Oxidative stress-induced senescence limits the persistence of chromosome 3p loss.**

**(A)** Hallmark gene set enrichment analysis (GSEA) comparing chromosome 3p loss cells to cells without chromosome 3p loss.

**(B)** GSEA plot showing enrichment for senescence-associated transcriptional program in chromosome 3p loss cells.

**(C)** Images of senescence-associated β-galactosidase (SA-β-gal) staining 5 days after transfection with the indicated sgRNAs.

**(D)** Quantification of SA-β-gal-positive cells shown in (C).

**(E-G)** GSEA plots showing enrichment of inflammatory response, oxidative phosphorylation (OXPHOS), and reactive oxygen species (ROS) pathways in chromosome 3p loss cells.

**(H)** Immunoblot confirmation of *RB1* knockout clones.

**(I)** Frequency of chromosome 3p loss in *RB1* WT and *RB1*^-/-^ cells at the indicated time points. Data are mean ± SEM from *n* = 2 independent experiments for *RB1* WT or *n* = 3 independent experiments for *RB1*^-/-^ from 53-120 metaphase spreads analyzed per condition.

**(J)** Immunoblot analysis of two independent cell populations overexpressing SOD3.

**(K)** Frequency of chromosome 3p loss in control and SOD3-overexpressing cells at the indicated time points. Data are mean ± SEM from *n* = 2 independent experiments from 51-167 metaphase spreads analyzed per condition.

**(L)** Frequency of whole-genome duplication (WGD) events in control, *RB1*^-/-^, or SOD3-overexpressing cells. Data were derived from 30, 100 and 70 metaphase spreads, respectively.

**Figure S7. Validation of dTAG-Flow for isolation of cells harboring chromosome 3p loss, related to Figure 3. (A-B)** Flow cytometry profiles of GFP-FKBP12^F36V^ (GFP-degron)-expressing cells with or without dTAGv1 treatment.

**(C)** Quantification of GFP+ cells following dTAGv1 treatment. Data are mean ± SEM of *n* = 5 independent experiments. Statistical analysis by unpaired t-test.

**(D)** Immunoblot analysis showing efficient degradation of GFP-degron following dTAGv1 treatment in *VHL* WT cells, partial degradation in *VHL*^+/-^ cells and incomplete degradation in *VHL*^-/-^ cells.

**(E)** Live-cell images of GFP-degron-expressing cells in the indicated VHL genetic backgrounds with or without dTAGv1 treatment.

**Figure S8. Chromosome 3p loss clones exhibit persistent stress-response adaptations.**

**(A)** Frequency of chromosome 3p loss cells recovered by dTAG-Flow enrichment at weeks 1 and 4. Data are mean ± SEM of *n* = 2 independent experiments from 37-155 metaphase spreads analyzed per condition.

**(B)** Interphase and metaphase FISH images of clone 1 showing the loss of the chromosome 3p arm. Scale bars, 5 µm (top) and 10 µm (bottom).

**(C)** Metaphase FISH image of clone 7 analyzed using chromosome 3 (green) and chromosome 5 (red) paint probes. Scale bar, 10 µm.

**(D)** Reduced expression of the major chromosome 3p-encoded ccRCC tumor suppressor genes (*VHL, PBRM1, BAP1,* and *SETD2*) in 3p loss clones as determined by bulk RNA sequencing. *TBP* and *SOX2* are included as non-chromosome 3p controls.

**(E)** Frequency of micronuclei in chromosome 3p loss clones. Data from 478, 510, 339, 385, 528, 457, 622, 503 and 753 interphase cells (left to right).

**(F)** Quantification of γH2AX intensity in WT RPTECs and chromosome 3p loss clones. Doxorubicin was used at the indicated concentrations as a control. au, arbitrary units. Statistical analysis by one-way ANOVA test with multiple comparisons.

**(G)** Metaphase FISH images showing telomeric sequences (telo) at terminal chromosome 3 breakpoints in chromosome 3p loss clones.

**(H)** Immunoblot analysis of SOD3 expression in WT RPTECs and chromosome 3p loss clones.

**(I)** Immunofluorescence images showing nuclear accumulation of SOD3 in clone 10. Scale bar, 20 µm.

**(J)** Quantification of nuclear SOD3 intensity shown in (I).

**Figure S9. Cytogenetic characterization of chromosome 3p loss clones, related to Figure 4.** Representative multiplex FISH karyotypes of WT RPTECs and chromosome 3p loss clones.

**Figure S10. Whole-genome sequencing reveals breakpoint-confined chromothripsis in clones with chromosome 3p loss, related to Figure 4.** Circos plots and genomic rearrangement profiles showing alterations across the indicated early-passage (18 days) RPTEC clones with chromosome 3p loss. Circos plot tracks depict, from outside to inside: (i) chromosome ideograms showing Giemsa banding data; (ii) copy number gains and losses shown in red and blue, respectively; and (iii) SVs depicted by colored lines corresponding to inversions (purple), deletions (blue), duplications (red), and translocations (black).

**Figure S11. Restoration of p53 signaling suppresses growth without reversing karyotypic evolution, related to Figure 4.**

**(A)** Immunoblot analysis showing doxycycline-induced expression of shRNA-resistant *TP53* in WT RPTECs and chromosome 3p loss clones.

**(B)** Clonogenic survival assays following induction of p53 expression.

**(C)** Quantification of clonogenic survival shown in (B).

**(D)** Soft agar colony formation assays following induction of p53 expression. Scale bar, 1 mm.

**(E)** Quantification of anchorage-independent growth shown in (D).

**(F)** Frequency of chromosome 5q gain in clone 7 before and after doxycycline-induced p53 expression. Data were derived from 250 metaphase spreads at 0 day and 154 metaphase spreads after 2 weeks of doxycycline induction.

**(G)** Frequency of chromosome 14q loss in clone 10 before and after doxycycline-induced p53 expression. Data were derived from 125 metaphase spreads at 0 day and 282 metaphase spreads after 2 weeks of doxycycline induction.

**Figure S12. Longitudinal genome evolution recapitulates recurrent ccRCC-associated rearrangements, related to Figures 4 and 5.** Circos plots and genomic rearrangement profiles showing alterations across the indicated late-passage (180 days) RPTEC clones with chromosome 3p loss. Circos plot tracks depict, from outside to inside: (i) chromosome ideograms showing Giemsa banding data; (ii) copy number gains and losses shown in red and blue, respectively; and (iii) SVs depicted by colored lines corresponding to inversions (purple), deletions (blue), duplications (red), and translocations (black).

**Figure S13. Chromosome 14q loss promotes metabolic adaptation and p62-dependent tumorigenic growth, related to Figure 6.**

**(A)** Immunoblot analysis of HIF1α expression in WT RPTECs and late-passage chromosome 3p loss clones.

**(B)** Immunoblot of two independent populations of WT RPTECs and chromosome 3p loss clones 7 and 10 overexpressing HIF1α.

**(C)** Clonogenic survival assays following overexpression of HIF1α.

**(D)** Soft agar colony formation assays following overexpression of HIF1α. Scale bar, 1 mm.

**(E)** Quantification of anchorage-independent growth shown in (D).

**(F-H)** Hallmark pathway enrichment analyses comparing WT RPTECs and late-passage clones 5, 7, and 10.

**(I)** Immunoblot analysis of p62, phospho-4E-BP1, and phospho-S6K1 following *SQSTM1* targeting with three independent sgRNAs.

**(J)** Soft agar colony formation assays following *SQSTM1* targeting. Scale bar, 1 mm.

**Figure S14. Late-evolved chromosome 3p loss clones generate metastatic renal tumors with ongoing genomic evolution, related to Figure 7. (B)** Representative primary tumors isolated from mice implanted with late-passage clone 10 cells.

**(C)** H&E staining and IHC for the indicated markers in primary tumors derived from clone 10. Scale bar, 100 µm.

**(D)** Immunoblot analysis showing retention of chromosome 3p-encoded tumor suppressor proteins, including pVHL, in tumor-derived cells.

**(E)** Multiplex FISH karyotypes of primary tumors and lung metastases showing continued chromosomal evolution during *in vivo* growth.

**(F)** Immunoblot analysis of p62 expression in parental and tumor-derived cells.

